# Trading off the cost of conflict against expected rewards

**DOI:** 10.1101/412809

**Authors:** Nura Sidarus, Stefano Palminteri, Valérian Chambon

**Affiliations:** Institut Jean Nicod, Département d’Études Cognitives, Ecole Normale Supérieure, EHESS, CNRS, PSL University, Paris, France; Laboratoire de Neurosciences Cognitives, Département d’Études Cognitives, Ecole Normale Supérieure, INSERM, PSL University, Paris, France

## Abstract

Value-based decision-making involves trading off the cost associated with an action against its expected reward. Research has shown that both physical and mental effort constitute such subjective costs, biasing choices away from effortful actions, and discounting the value of obtained rewards. Facing conflicts between competing action alternatives is considered aversive, as recruiting cognitive control to overcome conflict is effortful. Yet, it remains unclear whether conflict is also perceived as a cost in value-based decisions. The present study investigated this question by embedding irrelevant distractors (flanker arrows) within a reversal-learning task, with intermixed free and instructed trials. Results showed that participants learned to adapt their choices to maximize rewards, but were nevertheless biased to follow the suggestions of irrelevant distractors. Thus, the perceived cost of being in conflict with an external suggestion could sometimes trump internal value representations. By adapting computational models of reinforcement learning, we assessed the influence of conflict at both the decision and learning stages. Modelling the decision showed that conflict was avoided when evidence for either action alternative was weak, demonstrating that the cost of conflict was traded off against expected rewards. During the learning phase, we found that learning rates were reduced in instructed, relative to free, choices. Learning rates were further reduced by conflict between an instruction and subjective action values, whereas learning was not robustly influenced by conflict between one’s actions and external distractors. Our results show that the subjective cost of conflict factors into value-based decision-making, and highlights that different types of conflict may have different effects on learning about action outcomes.

## Introduction

Voluntary action depends on our capacity to learn how our actions relate to specific events in the external world, and use this knowledge to guide our decisions. Research on value-based decision-making has additionally revealed that the costs associated with specific actions, such as physical [1,2] or mental [3,4] effort, are weighed against their expected rewards [5,6]. In other words, when deciding whether to go out for dinner at a sushi or pizza restaurant, we consider not only how much we like either restaurant, but also how far we need to travel to reach them. Importantly, navigating the external world requires continuously monitoring our decisions, actions, and their consequences, to detect potential difficulties, or unexpected events, that may arise and adapt our behaviour accordingly. Returning to the dinner example, imagine you decide to go to the sushi restaurant but, as you step out of the house, you are faced with the smell of pizza from a new nearby restaurant. This will trigger a conflict between your previous plan to have sushi and the tempting smell of pizza, and may lead you to re-evaluate your decision.

Research on conflict monitoring has shown that detecting conflicts between competing alternatives leads to the recruitment of cognitive control resources [7,8]. Cognitive control serves to resolve conflict online by enhancing attention to task-relevant information [9,10], while sustained adjustments after conflict can reduce subsequent conflict effects [11–13]. As engaging cognitive control is effortful, experiencing conflict is typically considered aversive [14–16]. People tend to avoid high conflict tasks [17], and contexts associated with a high conflict probability, whether the conflict is triggered consciously [18,19] or unconsciously [20]. Many studies have also shown that our choices can be biased by both conscious [21,22] and unconscious [23–28] stimuli. Yet, in those studies, it was irrelevant whether participants chose one option or the other, as they had similar, or no, consequences. As choosing between options of similar value may itself constitute a type of conflict [28,29], following an external suggestion might even facilitate decision-making.

Remarkably, it remains unclear to which degree motivated, value-based *decisions* would be influenced by *irrelevant* conscious external stimuli. When choosing between options with different reward outcomes, we might expect people to only rely on learned, internal information, and ignore irrelevant external suggestions. Yet, as conflict resolution is perceived as effortful and aversive [30], and given the aforementioned research on effort costs [5], the subjective cost of conflict might be similarly traded off against expected rewards, predicting that external stimuli might still influence value-based decisions.

Independently of whether conflict costs factor into to our decision-making, experiencing conflict during a decision might also alter *learning* about action-outcomes associations, i.e. instrumental learning. In fact, the aversive nature of conflict has been shown to influence the processing of action outcomes. Conflicts can lead to a more negative evaluation of neutral stimuli [31,32], and a reduction in perceived control over action outcomes [33,34]. In line with findings on effort discounting [4,6], a study showed that response conflict may carry an implicit cost to obtained rewards [35]. Using the Simon task [36], Cavanagh and colleagues showed that participants preferred cue stimuli associated with rewards that followed non-conflicted trials, over stimuli associated with rewards that followed conflicted trials. Importantly, in the learning phase of that study, participants could not choose whether to avoid or experience conflict. Therefore, it remains unclear whether similar costs of conflict to learning would be found when participants can *choose* whether to experience conflict.

Finally, in addition to experiencing conflicts between internal and external information – akin to “pizza smell” example above, we can also experience conflicts between competing internal motivations – e.g. preferring sushi, but also wanting to please a friend who asks to have pizza. Interestingly, it has been shown that motivational conflicts, such as between Pavlovian biases and instrumental task requirements, can impair instrumental learning [37,38]. Thus, it is hard to learn to act to avoid punishments, as it goes against the Pavlovian tendency of withholding action to avoid punishments. Therefore, the precise nature of the conflict experienced – between internal vs. external information, or between competing internal motivations – may also be a relevant moderator of its effects on learning.

The present study aimed to investigate the following two key questions: a) whether value-based decisions could be influenced by irrelevant distractors, due to a subjective cost of conflict; b) whether experiencing conflict might influence instrumental learning. Additionally, we assessed the role of two potential moderators of how learning might be influenced by conflict: i) the type of conflict experienced – with external information, or between internal motivations; ii) having a choice in whether to experience conflict or avoid it. To test these questions, we embedded irrelevant distractors (flankers) within a reversal-learning task (Fig 1), with intermixed free and instructed trials. Participants had to continuously track whether left or right hand actions had a high or low reward probability (75/25%), and contingencies reversed unpredictably. As the same contingencies applied in free and instructed trials, participants were told to learn equally from the outcomes of both trial types, and that not complying with instructions would reduce their final earnings. Distractors could trigger conflict with an instructed action (e.g. >><>>) or with a freely chosen action (indicated by a bidirectional target), and might bias free choices. In this context, participants could adapt to conflict by focusing on the target and ignoring the distractors, while free choices additionally offered an opportunity for conflict avoidance. Comparing the influence of conflict on learning in free and instructed trials allowed us to assess the role of having choice in whether to act in conflict with an external suggestion. Furthermore, as instructions were equally likely to require making the high or low reward action, participants sometimes experienced conflict between two internal motivations: correctly following an instruction (e.g. left), and following their subjective value expectations about the best action (e.g. right).

**Fig 1.**
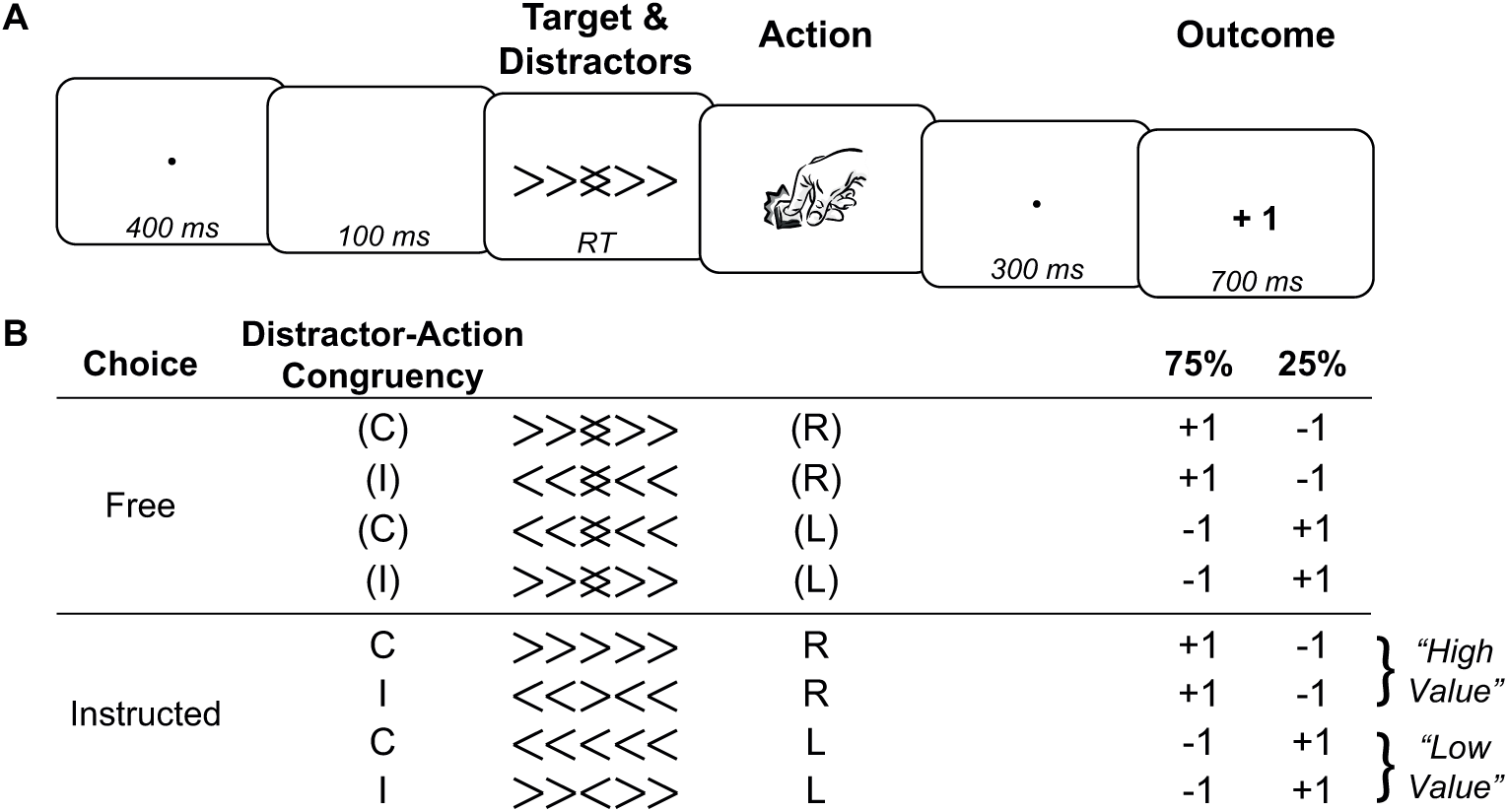
Task outline. **A.** Timeline of a trial. **B.** Task design and example mapping of actions to reward probabilities. Conflict between actions and external distractors is captured by the “distractor-action congruency” factor, where C=Congruent, and I=Incongruent. Conflict between instructions and subjective action values (model-based) is exemplified here. Assuming participants correctly learned the current contingency, right (R) would be the subjectively “high value” action (i.e. no conflict if instructed right), and left (L) would be the “low value” action (i.e. conflict if instructed left).

As brief glimpse of our findings, our results support our hypothesis that subjective conflict costs were traded off against off against expected. Moreover, learning was only influenced by conflict in instructed trials, when facing a motivational conflict between the instruction and subjective action values.

## Results

### Distractor effects on action

To verify that the distractors (i.e. flankers) elicited response conflict we analysed the effect of distractor congruency on different behavioural variables: reaction times, free choices and error rates.

#### Reaction Times

Mean reaction times (Fig 2A) were submitted to a repeated-measures ANOVA, as function of choice (free vs. instructed), and current distractor-action congruency (congruent vs. incongruent). This revealed a significant main effect of distractor-action congruency (*F*_1,19_ = 182.29, *p* < .001, 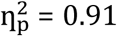), as well as a significant main effect of choice (*F*_1,19_ = 7.52, *p* = 0.01,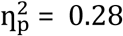), and a significant choice-by-congruency interaction (*F*_1,19_ = 59.78, *p* < .001,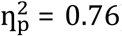). Post-hoc tests revealed that there was a significant congruency effect in both free and instructed trials (free: *t*_19_ =-10.60, *p* < .001, *d* = -2.38; instructed: *t*_19_ = -12.10, *p* < .001, *d* = -2.72), with slower RTs in incongruent than congruent trials. Additionally, for incongruent trials, RTs in instructed trials were significant slower than in free trials (*t*_19_ = -7.10, *p* < .001, *d* = -1.57), but there was no significant effect of choice in congruent trials (*t*_19_ = 0.78, *p* = .45, *d* = 0.17). These findings show that choosing to go against the distractors’ suggestion carried a cost to performance (i.e. slower RTs in free-incongruent than in free-congruent trials). Moreover, the cost of conflict to action selection was even greater in instructed trials (i.e. slowest RTs in instructed-incongruent trials). To correctly follow the instruction in incongruent trials, participants had to overcome conflict both at the level of the visual stimuli (to detect the target direction among opposing distractors), and of the response (as distractors and target triggered two competing responses).

**Fig 2.**
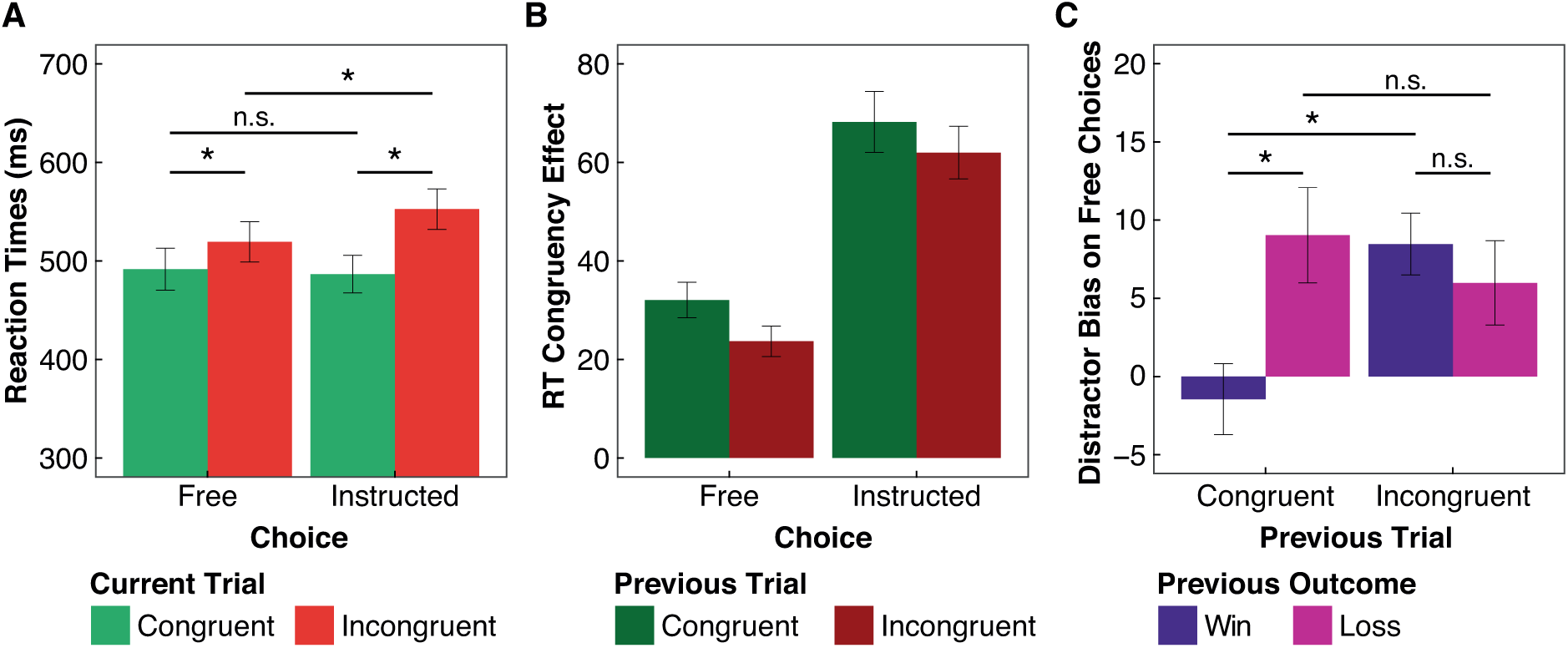
Influence of conflict at the decision stage. **A.** Average RTs as a function of choice and current trial distractor-action congruency. **B.** Average conflict effects on RTs (incongruent *minus* congruent) as a function of choice and previous trial congruency. **C.** Average distractor bias effect on free choices (percentage of congruent *minus* incongruent choices) as a function of previous trial congruency and outcome (win/loss).

#### Free Choices

. In free trials, participants were significantly biased towards choosing actions that were congruent with the direction of the distractors, than to make distractor-incongruent actions (congruent: 52.70% ±4.57; one sample t-test against 50% chance level: *t*_19_ = 2.65, *p* = .016, *d* = 0.84). Thus, participants generally tended to avoid conflict.

#### Errors

In instructed trials, participants made significantly more errors, i.e. not responding according to the target direction, when the distractors were incongruent with the target direction, than in congruent trials (congruent: 1.30% ±1.59; incongruent: 4.58% ±2.58; paired t-test: *t*_19_ = -5.78, *p* < .001, *d* = -1.29). This confirms that the distractors disrupted action selection, occasionally leading participants to make the wrong action.

### Conflict adaptation

To assess conflict adaptation, current trial congruency effects on RTs (incongruent *minus* congruent) were assessed as a function of choice (free vs. instructed) and previous trial congruency (congruent vs. incongruent; see Fig 2B). Repeated-measures ANOVA revealed a significant main effect of choice (*F*_1,19_ = 58.82, *p* < .001,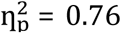), indicating that congruency effects were overall larger in instructed than in free trials, in line with the previous findings on RTs. Additionally, there was a significant main effect of previous trial congruency (*F*_1,19_ = 5.30, *p* = .03, 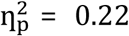), indicating conflict adaptation, as congruency effects on the current trial were reduced following incongruent trials, relative to following congruent trials. There was no significant interaction between the factors (*F*_1,19_ = 0.19, *p* = .67, 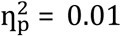), suggesting a similar conflict adaptation regardless of whether the current trial was free or instructed.

### Conflict avoidance

Next, we assessed how free choices were affected by both conflict and reward history (Fig 2C). We computed a “distractor bias” measure (percentage congruent *minus* incongruent free choices), which denoted the degree to which participants avoided conflict (by following the distractors’ suggestion). This “distractor bias” variable was submitted to a repeated-measures ANOVA with previous trial congruency and previous outcome (win vs. loss) as within-subject factors. Results showed a significant main effect of previous trial congruency (*F*_1,19_ = 5.49, *p* = .03,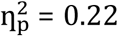), a significant main effect of previous outcome (*F*_1,19_ = 4.79, *p* = .04, 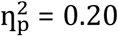), and a significant interaction between the two factors (*F*_1,19_ = 10.31, *p* = .005,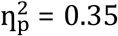). Post-hoc tests revealed that whereas the distractor bias was significantly larger after loss, than win, outcomes following congruent trials (*t*_19_ = -3.84, *p* = .001, *d* = -0.86), there was no effect of previous outcome following incongruent trials (*t*_19_ = 0.91, *p* = .37, *d* = 0.20). At the same time, the distractor bias was larger after incongruent, than congruent, trials following win outcomes (*t*_19_ = -5.01, *p* < .001, *d* = -1.12), whereas there was no effect of previous trial congruency following loss outcomes (*t*_19_ = 1.04, *p* = .31, *d* = 0.23). That is, if the previous trial was incongruent, or resulted in a loss, participants were more likely to be biased by the distractors’ suggestion, thus displaying greater conflict avoidance.

We hypothesise that recent conflict experience increases the importance of conflict avoidance, while losses might reduce the participant’s confidence in the best response. Thus, only when the last trial involved easy action selection and was rewarded (i.e. all “went well”) were participants *not* biased by the distractors (Fig 2C, one sample t-test of distractor bias in “previously congruent and win outcome” trials against 0 confirms there was no significant difference, *t*_19_ = -0.64, *p* = .53, *d* = -0.20; whereas distractor bias was larger than 0 in the remaining trial types, all *t*s > 2.22, *p*s < .039, *d* > 0.70).

### Learning Performance

To assess whether participants were able to accurately learn to make the best response, we calculated the percentage of high reward choices as a function of free trial number (i.e. ignoring instructed trials) relative to a reversal point. Given the choice bias identified above, we additionally considered how choices were affected by distractor-action congruency, and thus calculated the percentage of congruent or incongruent high reward choices, relative to the total number of free trials. The resulting learning curves (Fig 3A) show that participants successfully learned to maximise their rewards over time, matching objective reward probability level (75%, shown in Fig 3A divided by the 2 congruency conditions, i.e. 37.5%). In the free choice before a reversal, participants chose the high reward option on 76.13% ±0.09 of trials (one sample t-test against 50% chance level: *t*_19_ = 13.63, *p* < .001, *d* = 4.31). The learning curves additionally show that free choices were biased by the distractors, as participants were more likely to make congruent than incongruent choices, particularly at the start of learning episodes.

**Fig 3.**
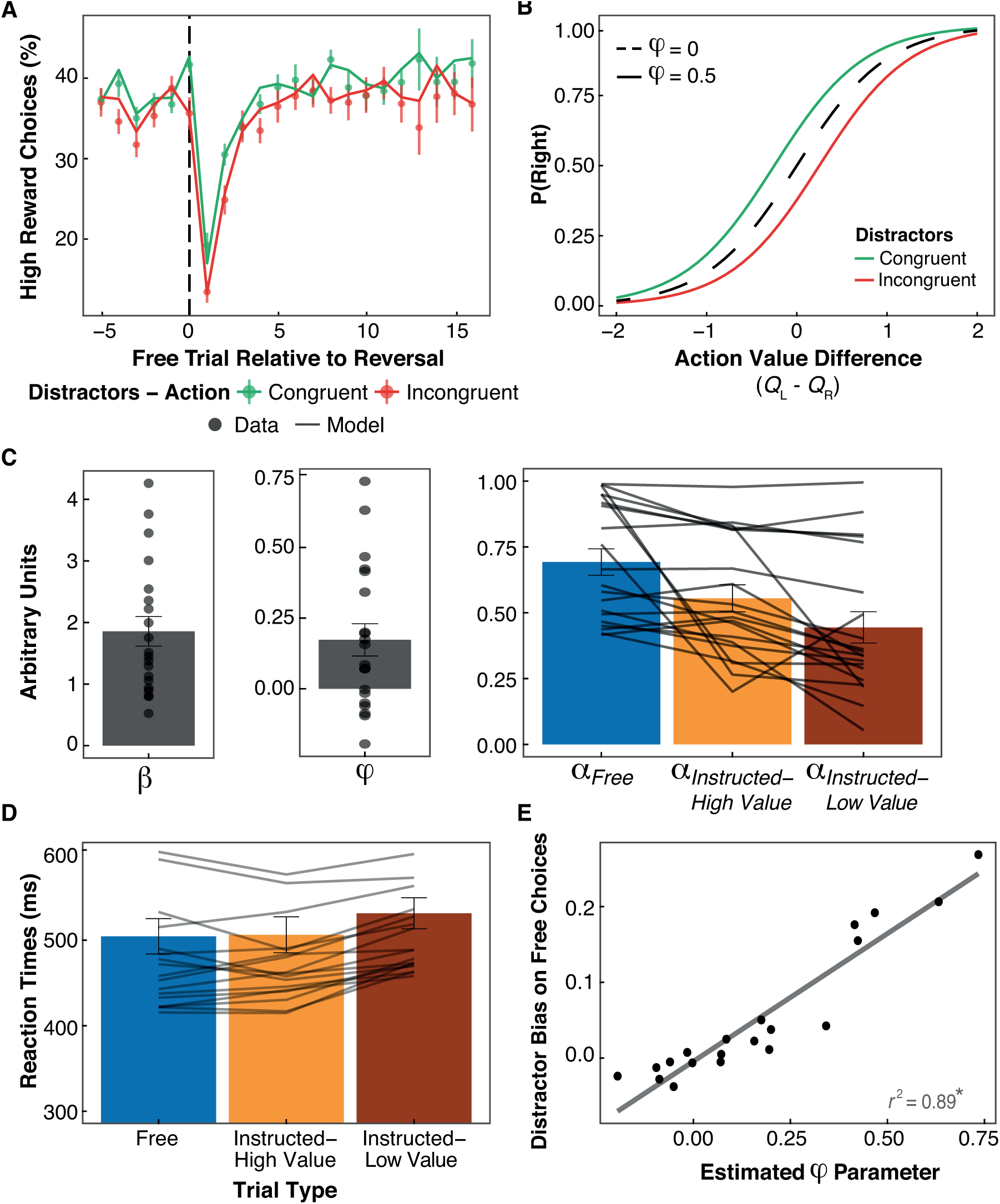
Model-based analyses of reinforcement learning (for wining model: m8). **A.** Learning curve (real data as points with standard errors, and model simulations as lines) displaying the percentage of high reward choices, and whether they were congruent or incongruent with the distractors, as a function of free trial number relative to a reversal point. **B.** Softmax decision rule for hypothetical distractor bias parameters (*φ* = 0 vs. *φ* = 0.5), and as a function of distractor-action congruency (for *φ* = 0.5). **C.** Average estimated parameters (as bars; with dots/lines representing each participant). **D.** Average RTs a function of choice, and whether instructions required making the subjectively high or low value action (based on simulated action (*Q*) values; lines represent each participant). **E.** Relation between the estimated distractor bias parameter (*φ*) and the observed distractor bias on free choices.

### Modelling the effects of conflict between action and distractors

Computational models of reinforcement learning [39,40] were adapted to better investigate our two research questions, i.e. the effects of conflict with external distractors on a) decisions, and b) learning (see Materials and Methods below for further details). At the decision stage, to capture the distractor bias on free choices described above, we adapted a standard softmax decision rule to include a distractor bias parameter, *φ*. This parameter can shift the softmax curve to increase the probability of making distractor-congruent choices (if *φ* > 0, see Fig 3B). As this distractor bias parameter (*φ*) captures the degree to which participants prefer to avoid conflict (by following the distractors), it thus serves as an implicit measure of the subjective cost of conflict. That is, participants with higher subjective conflict costs would show greater conflict avoidance, and hence a larger distractor bias parameter. At the learning stage, we used a standard *Q*-learning model to track how the expected action values (*Q*-values) are updated across trials. To test the effect of conflict between action and distractors on learning, we compared models with separate learning rates for congruent vs. incongruent trials, as well as a function of choice and choice-by-congruency interactions.

We initially compared a classic reinforcement learning model with one choice temperature and only one learning rate, with models that additionally included our distractor bias parameter, and potentially separate learning rates as a function of choice and congruency. Excluding the model that considered conflict between instructions and action values (described in the next section) from this initial model space allowed us to test whether there was *any* effect of action-distractor conflict on learning, rather than compare which type of conflict effects on learning might better fit the data. Model comparison (S1 Table) revealed that the winning model (m4) included the distractor bias (*φ*) parameter in the decision rule, and different learning rates as function of choice only (*xp = 0.97*). For this model, and in line with the observed choice bias towards distractor-congruent choices, we found that the estimated distractor bias parameter was significantly larger than 0 (average *φ* = 0.17 ± 0.25; one-sample t-test against 0: *t*_19_ = 3.05, *p* = .007, *d* = 0.97). Turning to the learning rates, a paired sample t-test revealed that learning rates were significantly higher in free than in instructed trials (*t*_19_ = 3.93, *p* < .001, *d* = 0.88).

Since the models with different learning rates as a function of congruency did not provide a significantly better fit to the data (considering the number of free parameters), we conclude that the present data did not reveal robust effects on learning of conflict between one’s action and irrelevant distractors (see S1 Text and S1 Fig for further analyses).

### Modelling conflict between instructions and subjective beliefs

To assess whether learning was influenced by motivational conflict between instructions and subjective action values, we considered an additional model (m8). Motivational conflict trials require resolving a conflict between two internal drives: choosing the most rewarding option, and following the instruction. Since errors in instructed trials reduced participants’ final earnings, they were still motivated to respond correctly, and follow the instruction. The extra model (m8) parses learning rates in instructed trials as a function of estimated action values (*Q*-values): if the instruction required the action favoured by the difference in action values, the trial was classed as “Instructed-High Value”, otherwise it was classed as “Instructed-Low Value”. As the previous model comparison showed an influence of choice on learning rates, this new model also allowed us to test whether the reduction in learning rates in instructed trials was specifically driven by trials in which participants had to follow an instruction that went against their subjective beliefs (i.e. instructed-low value).

Comparing this with all previously considered models allowed us to check the winning model provided the best fit to the data across the model space. Moreover, as the previous model comparison did not reveal robust effects of action-distractor conflict on learning, we did not additionally include models with both types of conflict. Model comparison across this extended model space (Table 1) showed that the new model (m8) provided a better fit to the data than all other models (*xp* = 0.97), including the previously winning model (m4). This confirmed that the conflict between the instruction and subjective action values had a robust influence on learning rates.

**Table 1.** Model comparison across the extended model space. *φ* refers to the distractor bias parameter added to the decision rule. α refers to the learning rates, split by conditions. C = Congruent, I = Incongruent, AIC = Akaike Information Criteria, SD = Standard Deviation.

| Models | AIC ± SD | Model Frequency | Exceedance Probability ($xp$) |
| --- | --- | --- | --- |
| m1: Standard RL [$\beta, \alpha$] | 723.3 ± 191.4 | 0.15 | 0.02 |
| m2: [$\beta, \varphi, \alpha$] | 714.3 ± 183.5 | 0.08 | 0.00 |
| m3: [$\beta, \varphi, \alpha_C \neq \alpha_I$] | 715.2 ± 184.0 | 0.04 | 0.00 |
| m4: [$\beta, \varphi, \alpha_{Free} \neq \alpha_{Instructed}$] | 704.7 ± 176.8 | 0.08 | 0.00 |
| m5: [$\beta, \varphi, \alpha_{Free\_C} \neq \alpha_{Free\_I} \neq \alpha_{Instructed}$] | 704.9 ± 178.4 | 0.06 | 0.00 |
| m6: [$\beta, \varphi, \alpha_{Free} \neq \alpha_{Instructed\_C} \neq \alpha_{Instructed\_I}$] | 705.2 ± 177.4 | 0.05 | 0.00 |
| m7: [$\beta, \varphi, \alpha_{Free\_C} \neq \alpha_{Free\_I} \neq \alpha_{Instructed\_C} \neq \alpha_{Instructed\_I}$] | 705.4 ± 178.8 | 0.10 | 0.00 |
| **m8: [$\beta, \varphi, \alpha_{Free} \neq \alpha_{Instructed\_High\ Value} \neq \alpha_{Instructed\_Low\ Value}$]** | **701.8 ± 177.8** | **0.44** | **0.97** |

To better understand how choice, and conflict between instructions and action values influenced learning, we submitted the estimated learning rates (Fig 3C) to a one-way repeated-measures ANOVA based on trial type (free, instructed-high value, instructed-low value). As expected, this showed a significant effect of trial type (*F*_1.84, 34.87_ = 13.72, *p* < .001, 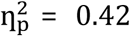). Bonferroni-corrected follow-up tests confirmed that conflict between the instruction and subjective beliefs led to a significant reduction in learning rates (instructed low vs. high value: *t*_19_ = 2.78, *p* = .036, *d* = 0.62). Moreover, the results showed that learning rates were significantly reduced by following instructions, even when the instruction required making what was believed to be the best action (free vs. instructed-high value: *t*_19_ = 2.70, *p* = .043, *d* = 0.60; free vs. instructed-low value: *t*_19_ = 4.89, *p* < .001, *d* = 1.09). This shows that the reduction in learning rates in instructed than free trials seen in the previous simpler model (m4) cannot be fully explained by trials in which participants were instructed to go against their beliefs.

### Relations between distractor bias parameter and behaviour

The estimated distractor bias in the new model (m8) was virtually identical to that estimated in previously winning model (average *φ* = 0.17 ± 0.26; one-sample t-test against 0: *t*_19_ = 3.03, *p* = .007, *d* = 0.96). To confirm that the estimated parameter was indeed related to participants’ choice bias, as expected, we assessed the correlation between the parameter estimates and the average distractor bias measure on free choices (percentage congruent *minus* incongruent). This confirmed a highly significant correlation (see Fig 3E; Pearson’s correlation: *r* = 0.94, *t*_18_ = 12.18, *p* < .001). [See S2 Text and S2 Fig for further correlations between the distractor bias parameter and behaviour, and S3 Fig from simulations demonstrating that the distractor bias did not robustly disrupt task performance.]

We further verified that our model could adequately replicate participants’ behaviour through model simulations. Based on the average estimated parameter values, we simulated data across the participants’ trial sequences. The resulting simulations can be observed in Fig 3A, and demonstrate that our model was able to replicate critical aspects of participants’ behaviour, such as the distractor bias on free choices (see S3 Text and S4 Fig for a demonstration that the *φ* parameter is essential to capturing this effect).

## Discussion

The present study investigated the influence of response conflict in value-based decision-making and learning. For this, we combined a flanker task, in which distracting stimuli could trigger response conflict, with a reversal-learning task, requiring the learning of action-outcome associations. To the best of our knowledge, this is the first study to show that even motivated – value-based – choices can be biased by irrelevant external information. Our findings, summarised in Fig 4, suggest that this bias results from a trade-off between the expected value of a given action and the cost of triggering response conflict, by going against an external suggestion. At the learning stage, we found that participants updated their value representations less when they had to follow instructions than when they could freely choose what to do. Learning was further reduced when participants had to follow instructions that went against their subjective beliefs. Thus, we found that learning was influenced by conflict between instructions and subjective action values, but did not find robust evidence that learning was influenced by conflict between one’s actions and external distractors.

**Fig 4.**
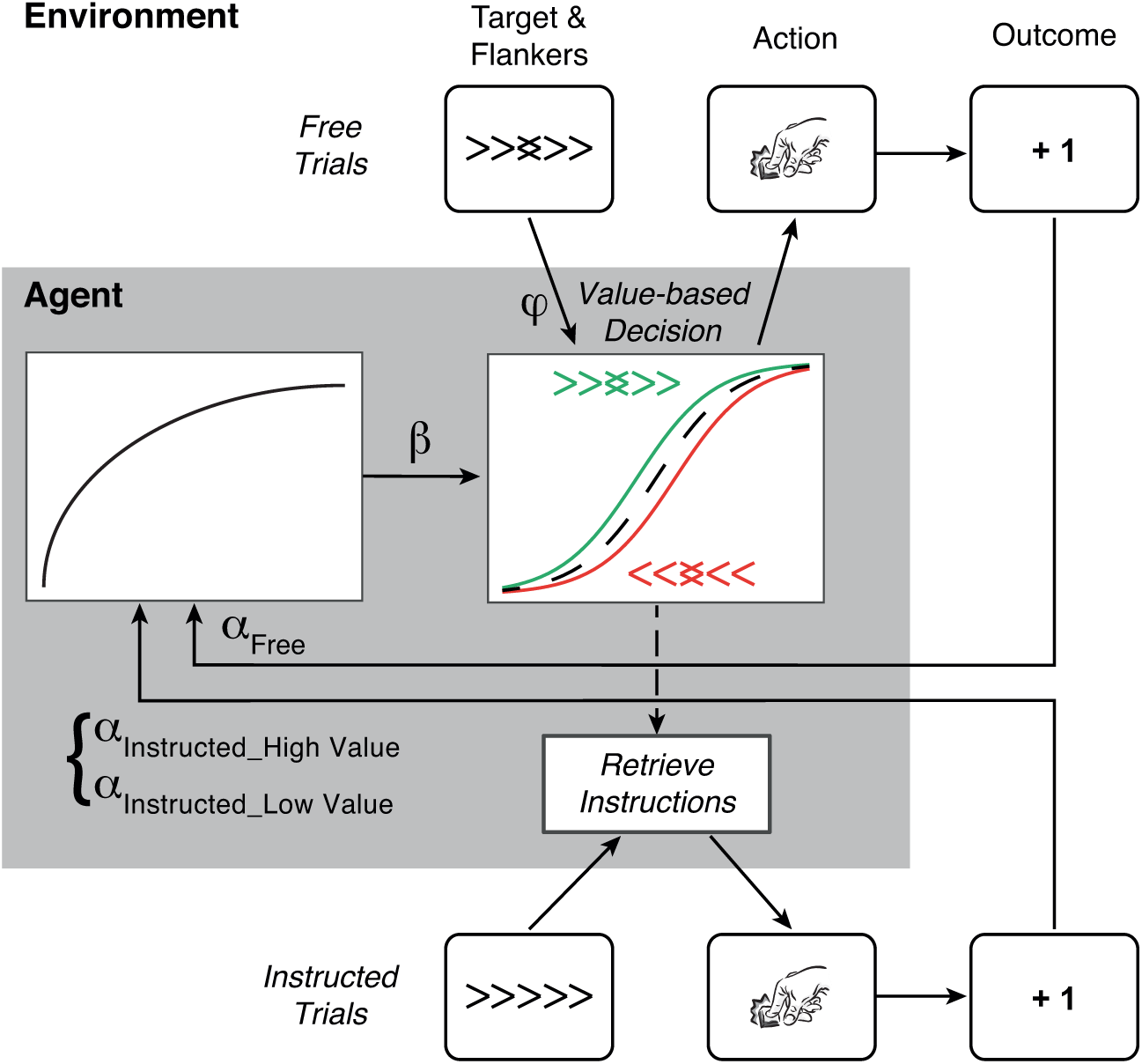
Schematic representation of the winning computational model. In Free trials, participants make value-based decisions by integrating internal information about action values (as a function of the inverse temperature parameter, *β*) with the external suggestion of the distractors (as a function of the distractor bias parameter, *φ*). The distractor bias parameter, *φ*, captures participants’ tendency to avoid conflict, by increasing the likelihood that participants will choose actions congruent with the distractors, especially when the difference in action values is small (i.e. larger gap between green/red lines in the middle of the graph, see also **Fig 3**B). The outcomes of those freely chosen actions can then be used to update action values (as a function of the learning rate, *α*_*Free*_). In instructed trials, participants must retrieve the instructions of following the target direction, but the accumulated action value information may partially interfere with their responses (influence on RTs seen in **Fig 3**D). Moreover, action outcomes were used to update actions values differently depending on whether the instruction required making the subjectively high vs. low value action (*α*_*Instructed_High*_ Value vs *α*_*Instructed_Low Value*_ respectively, based on the previously accumulated values).

### Influences on decision-making

At the decision stage, we found that decisions guided by internal value representations could be biased by external information. When free to choose what to do, participants typically chose the most rewarding action, but were also more likely to choose an action that was congruent with the suggestion of distracting stimuli, than to choose the opposite action. Moreover, choosing to go against the distractors’ suggestion was associated with slower RTs than following the distractors’ suggestion (Fig 2A). This confirms that response conflict was triggered by distracting stimuli that were incongruent with participants’ choices, resulting in a cost to action selection. These findings are consistent with previous studies showing conflict costs associated with choosing to go against conscious [22] and unconscious [25,27,28,41] distracting stimuli.

In instructed trials, when participants had to follow the target’s direction, RTs were also slower if the distractors were incongruent with the target, and the required action, relative to congruent distractors (Fig 2A). Results further showed that RTs were slower in instructed than in free trials when the distractors were incongruent with the executed action, whereas RTs did not differ between free and instructed trials with congruent distractors. Therefore, the cost of conflict in instructed trials was larger than in free trials. This added difficulty may reflect the fact that instructed-incongruent trials involved both a conflict at the perceptual level, as the target differed from the surrounding distractors (e.g. <<><<), as well as at a response level, between the competing responses triggered by the target and distractor stimuli. Such an interaction between choice and congruency has been previously report with the flanker task [22].

Considering how actions were affected by recent conflict experience revealed the well documented conflict adaptation effect [12,13], of reduced conflict effects on RTs following conflict trials (Fig 2B). Although conflict effects were generally smaller in free trials (as discussed above), the reduction in the cost of conflict on RTs *following* conflict was similar in free and instructed trials. This is consistent with previous conflict triggering behavioural adjustments, such as greater attention to the middle target and/or suppression of distractors, thus similarly reducing the impact of conflicting distractors on RTs across free and instructed trials. To the best of our knowledge, this is the first study to show such conflict adaptation effects in the context of intermixed free and instructed trials, and within a reinforcement learning task. These findings demonstrate the generalizability of such adaptation processes, which can be effective even in the presence of concurrent task demands (i.e. learning about action-reward contingencies).

The observed bias in participants’ choices to follow the distractors’ suggestions may serve as another conflict adaptation mechanism, in order to avoid experiencing future conflicts [8,42]. This account is supported by the observation that choices were *more* biased by distractors following conflict trials (Fig 2C), suggesting that recent conflict experience may highlight the aversive nature of conflict, and drive conflict avoidance. It is worth emphasising that the probability of left and right distractors was equated, hence the distractors were equally likely to suggest the high or low reward action. Participants were made aware of that, and instructed to ignore the distractors. As we found that participants could learn to maximise their earnings, the distractor-related choice bias did not impair participants’ performance (see S2 Text and S2 and S3 Figs for further demonstrations that the distractor bias did not carry a relevant cost to performance). In fact, the observed learning curves (Fig 3A) suggest that the distractor bias was larger after reversals, when participants were more uncertain about which was the best action.

Our computational model (Fig 4), captured this effect by adding a “distractor bias” parameter, *φ*, to the decision rule (Fig 3E shows a clear correlation between the observed choice bias and our bias parameter). Although, theoretically, even small value differences could determine a decision, reinforcement learning models incorporate a degree of stochasticity in the decision rule through the temperature parameter, which is proportional to the value difference. In our model, the distractor bias parameter shifts the decision rule to increase the probability of distractor-congruent choices (if *φ* > 0, as observed), simultaneously reducing the probability of distractor-incongruent choices (Fig 3B). The distractor bias parameter could thus be interpreted as reflecting the subjective cost of conflict, as a larger bias value would require a larger expected reward to increase the probability of acting in conflict with the distractors. These findings are consistent with recent work demonstrating that the costs of exerting cognitive control are weighed against expected returns [43,44], similarly to other types of mental [3] and physical [6] effort discounting. Moreover, as stochasticity is greater for smaller value differences, our model naturally entails a greater influence of distractors on decisions with greater uncertainty about the best action (see Fig 3B). It could be argued that the distractors could have *facilitated* the decision. As deciding between similarly valued options may be seen as a type of conflict in itself [28,35], the distractors’ suggestion could help break the tie between similar alternatives. The present results do not allow us to unequivocally arbitrate between the conflict avoidance vs. facilitation accounts. To avoid lengthening the experiment, the present study did not include trials with neutral distractors. Future studies with neutral distractors, and modelling differential facilitation and conflict bias parameters, could potentially dissociate an increased probability of distractor-congruent actions (facilitation) from a reduced probability of distractor-incongruent actions (conflict avoidance), relative to the neutral condition. Yet, these two accounts may not necessarily be incompatible.

From the perspective of decision-making as involving the accumulation of evidence for a response [45,46], both internal and external sources of information are integrated over time, until a decision bound is reached. As time-pressure was present in our study, delaying the response too much was counterproductive. Speed-accuracy trade-offs have been shown to influence evidence accumulation [47]. High expected values would lead to faster decisions than low values [29], and potentially to preparing the high value response *before* the trial. Such advanced preparation is supported by our findings that RTs were similar in free trials and when an instruction required making the subjectively best choice (Fig 3D), as both allowed participants to go ahead with the prepared action. In contrast, the presence of incongruent distractors would provide evidence for the alternative (low value) response, thus delaying the decision (i.e. slower RTs in incongruent than congruent trials, Fig 2A). When the initial evidence bias was high (given the expected value), recruiting control to suppress the effect of the distractors would be justified by enabling a quicker decision, since the accumulated evidence would remain closer to the decision bound of the high value option. Conversely, when the initial value difference was small, suppressing the distractors would return the accumulated evidence near the starting point, further delaying the response. In such cases, engaging cognitive control to suppress the distractors might carry the additional opportunity cost of foregoing any reward at all (as “too slow” responses constituted an error). The resulting facilitation of the decision by following the distractors’ suggestion would thus serve to avoid unnecessary/unjustified conflict. Future studies employing a combination of drift-diffusion and reinforcement learning models [29], together with time-sensitive neuroimaging techniques (e.g. M/EEG), might yield important insights into the process of integrating multiple sources of information for decision-making, and dealing with potential conflicts between them.

### Influences on learning

Results showed that learning rates were higher in free choices than in instructed trials (Fig 3C). This suggests we might learn more about the consequences of actions that are driven by our own intentions and motivations. A greater sensitivity to rewards obtained through one’s choices over passively received rewards has been previously shown [48], and having a choice in what to do may itself be rewarding [49,50]. Moreover, an “illusion of control” has been demonstrated [49,51], wherein more favourable outcomes are expected for one’s choices, over when one has no choice. An increased value and reward expectation for one’s choices, combined with a tendency for learning more from positive feedback [52,53], may thus boost learning from free choices over instructed actions.

Research on learning and memory has shown improvements when people are allowed to decide how, or which items, to study, relative to not having a choice [54,55]. Being able to choose what to do, also referred to as self-directed learning, allows for more efficient deployment of resources to relevant information gathering [56], such as testing relevant hypotheses (i.e. is this really the best action?), in turn improving learning. When following instructions, one is exposed to information that may not seem particularly informative (if the other action yields a reward, that could just reflect a low probability outcome, rather than a reversal in contingencies). Furthermore, different neural mechanisms have been linked to learning from one’s choices relative to the choices of others [57,58], even when similar learning performance is demonstrated in a separate, post-learning, test. As our analyses focus on the dynamics of value updating in a frequently changing environment, they may emphasise differences in learning mechanisms from free vs. instructed actions.

Turning to the effects of conflict on reinforcement learning, our results suggest that the type of conflict experienced is important. Our task was primarily designed to induce conflict between one’s actions and external distracting information (i.e. flankers). Yet, in instructed trials, another type of conflict could be elicited between the instruction and subjective action values – i.e. what participants *had* to do vs. what they *wanted* to do. We found that both types of conflict disrupted action selection. Incongruent distractors led to slower RTs than congruent distractors, and triggered sequential conflict adaptation in RTs and choices (Fig 2). Conflict between instructions and subjective values also led to slower RTs, relative to when the instruction required the high value action or free choices (Fig 3D). Yet, whereas conflict between instructions vs. subjective values (“instructed-low value” trials) led to a reduction in learning rates (Fig 3C), relative to no conflict (“instructed-high value” trials), we did not find robust evidence that conflict between actions vs. distractors modulated learning rates.

We suggest conflict between instructions and subjective values constitutes a type of motivational conflict. In those trials, participants were faced with two internal motivations competing to guide action selection: using subjective value information to make the best decision, vs. correctly following the instructions, to avoid losing potential rewards. We found that when participants had to suppress the drive to make what they perceived to be the best action, in order to correctly follow a subjectively “bad” instruction, they learned less from the observed outcomes. Thus, the conflict experienced during action selection seemed to devalue the action outcome. Such costs of motivational conflict to learning are consistent with previous studies involving conflict between Pavlovian tendencies and instrumental task requirements [37,38].

Conflict between one’s actions and external distractors might constitute a quite different type of conflict from motivational conflict. Although similar neural markers of conflict monitoring (i.e. frontal theta) have been found with the flanker task [59] and motivational conflicts [60], the cognitive control resources needed to resolve these two types of conflict might differ, and hence carry different subjective costs. Results showed that modelling different learning rates a function of conflict between action and external distractors (i.e. distractors) did not sufficiently improve model fit to justify the extra model complexity (see S1 Text and S1 Fig for further consideration of the effects of this type of conflict on learning rates). It remains possible that the current design, or our sample size, limited our ability to detect an influence of action vs. distractor conflict on learning. Additionally, the binary categorisation of trials into conflicted/non-conflicted (i.e. incongruent vs. congruent) may not have been sufficiently sensitive to trial-by-trial variations in the degree of conflict experienced.

Alternatively, conflict triggered by irrelevant external stimuli may lead to more targeted conflict resolution mechanisms, focused on suppressing irrelevant information, in turn reducing its impact on outcome evaluation. Such conflict adaptation processes may be more difficult to engage, or less efficient, in the context of motivational conflicts. Whereas participants could systematically ignore the distractors in our task, they had to constantly switch between using subjective values to guide their decisions and following the task instructions (i.e. target direction). Therefore, in addition to the type of conflict, the possibility, or capacity, for adaptation to conflict might be a relevant modulator of the effect of conflict on learning. Similarly, being able to choose *whether* to trigger conflict may also be relevant. Arguably, the absence of any effects of conflict on learning in free trials seen here might provide some initial support for a moderating role of choosing whether to trigger or avoid conflict (see S1 Text for further consideration of this hypothesis, namely as account for different results from [35]).

## Conclusions

Our findings suggest that decision-making involves trade-offs between the expected value of a given course of action and the potential cognitive control costs incurred by that action. Unless there is a sufficiently good reason to trigger it, e.g. expecting a reward, conflict is typically avoided. While experiencing conflict can sometimes influence subsequent processes, such as outcome evaluation, our results suggest that instrumental learning may not always be affected. We speculate that the effect of conflict on learning may be moderated by the type of conflict experienced, i.e. between competing internal drives or between internal vs. external information, as well as by the potential for conflict adaptation and avoidance.

## Materials and Methods

### Participants

Twenty participants completed the study (10 females, mean age = 25.80 ±4.34). One participant had been recruited and completed the first session, but due to problems scheduling the second session, was excluded. The study was conducted in accordance with the declaration of Helsinki (1964, revised 2013), and was approved by a local ethics committee. Participants gave written informed consent to participate in the study, and were told they would receive 15€ payment, and up to 5€ extra based on their performance, per session (~1.5 h). In fact, every participant received 20€ per session. All were right-handed, with normal or corrected-to-normal vision, did not suffer from colour blindness, and reported having no history of psychiatric or neurological disorders.

### Materials

Participants were seated approximately 50 cm from a computer screen. The experiment was programmed and stimuli delivered with Psychophysics Toolbox v3 [61–63], running on Matlab (MATLAB 8.1, The MathWorks Inc., Natick, MA, 2013). Stimuli were presented in black on a half grey background. A fixation dot was presented subtending 0.26° visual angle. Target and flanker stimuli consisted of left or right pointing arrows, subtending 0.6° visual angle, with a spacing of 0.1° between arrows. Participants responded by pressing one of two keys on a keyboard. Outcome stimuli consisted of “+1” or “-1”, presented in 54 points Arial font. The error cross subtended 1° visual angle.

### Task

The reversal-learning task (Fig 1) required participants to continuously learn the reward probabilities (75%/25%) associated with left vs. right hand actions, and adapt their choices accordingly. For example, right hand actions might have a 75% probability of yielding a reward (+1 point), whereas left hand actions would yield a reward in only 25% of trials. The remaining trials were associated with a loss (−1 point). This task was combined with a flanker task, such that participants responded to a target arrow, which appeared surrounded by irrelevant distractor (i.e. flankers). Each trial started with a 400 ms fixation dot, followed by a 100 ms blank screen. The target and distractor array was displayed until a response was made, or up to 1.2 s. In free trials, the middle arrow consisted of two overlapping left/right pointing arrows, indicating that participants were free to choose which action to make. Trials were classed as *congruent* if participants chose the action that corresponded to the direction of the distractors, and as *incongruent* if participants chose the opposite action to the distractors. In instructed trials, the target arrow consisted of a left or right pointing arrow, and participants had to respond according to its direction. Distractors could be congruent or incongruent with the target direction. If participants responded correctly, after a brief interval of 300 ms, the reward outcome (+1/-1) was displayed for 700 ms. The inter-trial interval varied randomly between 0.8-1.2 s. If participants made the wrong action in instructed trials, or did not respond within 1.2 s, an error cross was immediately displayed for 700 ms. To ensure participants responded correctly in instructed trials, they were informed that errors would lead to a deduction from the final earnings they obtained in the learning task.

After an unpredictable number of trials, the mapping of action to reward probabilities was reversed. After a reversal, the best-rewarded response (e.g. right) became the least-rewarded response, and vice-versa. The length of reversal episodes, i.e. number of trials before a reversal, followed a pseudo-gaussian distribution ([8, 8, 16, 16, 16, 24, 24, 24, 32, 32]). To ensure that all conditions (free vs. instructed, congruent vs. incongruent) were adequately counterbalanced within each episode, the same number of free and instructed trials was included. In instructed trials, we ensured equal numbers of left/right congruent/incongruent trials. As we could not control congruency in free trials, we presented equal numbers of left and right pointing distractor. Outcome probability was equated across congruency conditions for the instructed trials. Across the experiment, we counterbalanced the condition of the first trial in an episode (free/instructed, left/right distractors, and left/right instructed actions). The distribution of episode lengths and type of trial at the start of the episode was pseudo-randomised, such that all combinations of lengths and types were randomised before being repeated. We additionally ensured that the same type/length (or trial type in randomising trials) was not repeated more than 3 times in a row. Participants completed a total of 1600 trials, across 2 sessions (separated by ~5 days, range: 1-8). Breaks were introduced approximately every 15 mins, for 10 s. Participants were instructed that the breaks were independent of changes in reward probabilities.

Before the main experiment, participants completed some training blocks. During training, the reward probabilities were made more easily distinguishable (87.5% vs. 12.5%). Additionally, in the first training block, the best action was cued by displaying the corresponding target arrow in green (when the option was available). This served to help participants understand the probabilistic nature of the rewards, and track the reversal of the reward probabilities. In the second training block, the best action was no longer cued, thus all arrow stimuli were presented in black, as in the main experiment.

### Behavioural analyses

Trials without a response within 1.2 s were excluded (free: 0.90% ±2.08; instructed: 0.71% ±1.83). In instructed trials, the percentage of error trials was analysed as a function of distractor-target congruency, and errors were excluded from further analyses. Other statistical tests are described in the results section.

### Computational models

We fitted the data with a standard *Q*-learning model [39,40]. The model estimates the expected values (*Q*-values) of the two possible actions (left vs. right hand). The *Q*-values were set to 0 before each learning session, corresponding to the *a priori* expectation of a 50% chance of winning 1 point, plus a 50% chance of losing 1 point. After each trial *t*, the value of chosen option was updated according to the following rule:

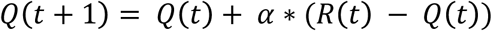

where *R*(*t*) was the reward obtained for the chosen option at trial *t*, and *α* referred to the learning rate parameter.

A softmax rule was adapted to estimate the probability of selecting a particular option (e.g. right) as a sigmoid function of the difference between the net values of left and right options, with a temperature parameter *β*, which captures choice stochasticity. To capture the influence of the distractors at the decision-making stage, we added a free parameter *φ*, which biased the choice depending on the relation between the distractors and that option at trial *t* as follows:

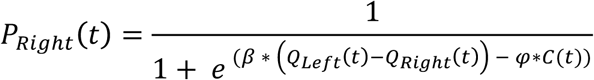

where *C*(*t*) = 1 if distractors were congruent with that option (i.e. point to the right), and *C*(*t*) = –1 if distractors were incongruent with the option. Thus, the estimated parameter *φ* > 1 indicates that distractors biased choice toward the congruent option; *φ* < 1 indicates that distractors biased choice toward the incongruent option; or *φ* = 0 indicates that the distractors had no influence on choice.

As the model-free analyses revealed that free choices were significantly biased by the distractors, we considered it critical to capture this influence at the decision stage in our models. Therefore, apart from considering the simplest, standard reinforcement learning model (with only two free parameters [*β, α*]), the remaining models in our model space included the *φ* parameter in the decision rule (see S3 Text and S4 Fig for evidence that this parameter is needed to adequately capture participants’ behaviour). To assess the potential influence of the choice and congruency manipulations on learning, we considered models that varied in the number of learning rates, from a single learning rate, to different learning rates as a function of choice, congruency, and their interaction. In supplementary simulations analyses we validated our models and parameter optimisation procedure to ensure we could dissociate effects of conflict at the decision vs. learning stages (see S4 Text and S5 Fig).

In an extra model (m8), we estimated separate learning rates as a function of choice, and as function of subjective beliefs for instructed trials. That is, we used the estimated difference in action values (Δ*Q = Q*_*L*_ – *Q*_*R*_) to split trials with instructions to make the (subjectively) high vs. low value action. For example, if participants followed an instruction to go right:

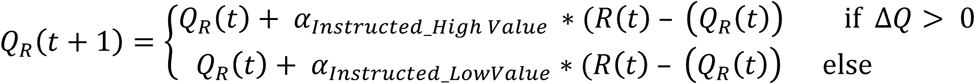

### Parameter optimisation and model comparison

Model parameters were optimised by minimising the negative log-likelihood of the data, given the parameters settings (using Matlab fmincon function, ranges: 0 < *β* < +Infinite, - Infinite < *φ* < +Infinite, and 0 < *α*_*n*_ < 1). To compare models fits while accounting for the model complexity of adding extra free parameters, we calculated Aikake Information Criteria (AIC) based on the negative log-likelihoods for each participant, and each model, as follows:

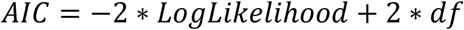

where *df* refers to the number of free parameters. In this study we opted for the AIC over the BIC, because the latter criterion tended to over-penalized complex models.

AIC values were then used as an approximation to the log model evidence [64], and models were treated as a random variable in a group-level variational Bayes analysis for model selection (using the “VBA toolbox” [65]). This approach allows for the estimation of the expected model frequency, and exceedance probability of each model, within the model space and given the data from all participants [66]. Expected model frequency quantifies the posterior probability of the model, i.e. the likelihood that the model generated the data of a random subject in the population. The exceedance probability (*xp*) quantifies the probability that a given model fits the data better than all other models in the set, i.e. has the highest expected frequency.

Following previous work [67,68], we conducted an additional optimisation procedure that minimised the logarithm of the Laplace approximation to the model evidence (or log posterior probability, LPP). This approach avoids degenerate parameters estimates as it includes priors over the parameters (Gamma(1.2,5) for *β*; Normal(0,1) for *φ*; and Beta(1.1,1.1) for *α*_*n*_). Further analyses (and figures) are based on the parameters estimated though this procedure.

## Data availability

Data is available at Open Science Framework: https://osf.io/cd62a/

## Author contributions

Conceptualization: NS SP VC.

Data curation: NS.

Formal Analysis: NS.

Funding Acquisition: NS VC.

Investigation: NS.

Methodology: NS SP VC.

Project Administration: NS.

Resources: VC.

Software: NS SP VC.

Supervision: VC.

Validation: NS SP VC.

Visualization: NS.

Writing – Original Draft Preparation: NS.

Writing – Review … Editing: NS SP VC.

## Supporting Information

**Table S1.** Reduced model space comparison results. *φ* refers to the distractor bias parameter added to the decision rule. α refers to the learning rates, split by conditions. Winning model (m4) highlighted in bold. C = Congruent, I = Incongruent. AIC = Akaike Information Criteria, SD = Standard Deviation.

| Models | AIC ± SD | Model Frequency | Exceedance Probability ($xp$) |
| --- | --- | --- | --- |
| m1: Standard RL [$\beta, \alpha$] | 723.3 ± 191.4 | 0.14 | 0.02 |
| m2: [$\beta, \varphi, \alpha$] | 714.3 ± 183.5 | 0.10 | 0.00 |
| m3: [$\beta, \varphi, \alpha_C \neq \alpha_I$] | 715.2 ± 184.0 | 0.06 | 0.00 |
| **m4: [$\beta, \varphi, \alpha_{Free} \neq \alpha_{Instructed}$]** | **704.7 ± 176.8** | **0.43** | **0.97** |
| m5: [$\beta, \varphi, \alpha_{Free_C} \neq \alpha_{Free_I} \neq \alpha_{Instructed}$] | 704.9 ± 178.4 | 0.09 | 0.00 |
| m6: [$\beta, \varphi, \alpha_{Free} \neq \alpha_{Instructed_C} \neq \alpha_{Instructed_I}$] | 705.2 ± 177.4 | 0.06 | 0.00 |
| m7: [$\beta, \varphi, \alpha_{Free_C} \neq \alpha_{Free_I} \neq \alpha_{Instructed_C} \neq \alpha_{Instructed_I}$] | 705.4 ± 178.8 | 0.12 | 0.01 |

**S1 Text. Effect of action vs. distractor conflict on learning**

For the sake of full disclosure and transparency, we present here a summary of the estimated parameters obtained by m7 – a model with separate learning rates as function of the interaction between choice and distractor-action congruency, plus the distractor bias parameter in the decision rule. Since this model did not win over the others in the models comparison, due to the extra model complexity, these results are only suggestive, and should be interpreted with care. We still thought it important to share these results, since the only weak evidence we found for an effect on action vs. distractor conflict on learning would be a benefit to learning. This goes against the hypothesis that conflict generally carries a cost to learning, due to its aversive nature, and is the opposite of what we found for conflict between instructions and subjective values. Finally, although the hypotheses discussed here remain speculative, they may offer relevant ideas for future research.

As in the other models, the estimated distractor bias parameter was significantly different from 0 (average *φ* = 0.17 ± 0.25, *t*_19_ = 3.05, p = .007, d = 0.96). The estimated learning rates (see S1 Fig) were submitted to a repeated-measures ANOVA (choice: free vs. instructed; distractor-action congruency: congruent vs. incongruent). This showed a significant main effect of choice (*F*_1, 19_ = 15.56, *p* < .001, 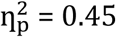), as learning rates were lower in instructed than free choices. It additionally showed a significant main effect of distractor-action congruency (*F*_1, 19_ = 14.98, *p* = .001, 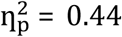), with higher learning rates in incongruent than congruent trials. Yet, these main effects were qualified by a significant choice-by-congruency interaction (*F*_1, 19_ = 5.94, *p* = .02, 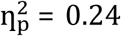). Post-hoc tests revealed that learning rates were significantly higher in incongruent than congruent trials in instructed trials (*t*_19_ = -4.85, *p* < .001, *d* = -1.08), but there was no significant effect of congruency in free trials (*t*_19_ = -0.12, *p* = .906, *d* = -0.03). Moreover, learning rates were always lower in instructed than in free trials (free-congruent vs. instructed-congruent: *t*_19_ = 4.28, *p* < .001, *d* = 0.96; free-incongruent vs. instructed-incongruent: *t*_19_ = 3.14, *p* = .005, *d* = 0.70).

These results suggest that in instructed trials, the conflict triggered by incongruent distractors and targets might have led to an increase in learning rates, relative to no conflict. Yet, in free trials, action vs. distractor conflict did not influence learning. Recall that we found larger RTs cost in instructed than free trials due to conflict (in main results). We hypothesised this was related to increased conflict costs associated with resolving both perceptual conflict (target vs. distractors), and at a response level (simultaneous activation of both responses), whereas in free choices, there was only conflict at the response level. This combined perceptual and response conflict in instructed trials may have required the deployment of more attentional resources to focus on the relevant stimuli, than in free trials. This might in turn result in enhanced attention at the time of the outcome, and thus in higher learning rates in incongruent than congruent trials.

It could have been argued that the absence of conflict effects on learning rates in free relative to instructed trials would be linked to differences in the effect sizes. RTs were slower in incongruent than congruent trials by an average of 66 ms in instructed trials, but only around 28 ms in free trials. This reduced effect might have thus been too weak to influence learning. However, the effects we found on learning rates due to conflict between instructions and subjective values (in m8) were associated with around 25 ms conflict costs on RTs (instructed low *minus* high value), similar to the cost of distractor-action conflict in free trials. Hence, the relatively smaller RTs costs in free trials cannot explain the absence of an effect on learning.

Therefore, the absence of effects of conflict on learning in free trials might rather be due to conflict being dealt with differently. Although free choices were still disrupted by incongruent distractors (evidenced by slower RTs, Fig 2A), such choices were likely driven by large differences in action value (as implied by our model). We speculate that such chosen conflict might be subjectively experienced as different from imposed (or unavoidable) conflict. The extra effort might seem “justified” by the expected action values, rendering it less aversive, and cancelling out potential conflict costs on learning [69]. Furthermore, as participants might generally devote more attention to the task in free trials, due to a greater perceived relevance of information (as seen in the choice effect on learning rates), the attention at the time of the outcome might not be further modulated. In contrast, if participants are less engaged in the task in instructed trials, but are then obliged to pay attention to the task to successfully resolve conflict, this enhanced attention may then improve outcome processing, relative to the instructed-congruent trials.

We further suggest that having a choice in whether to experience conflict may partially explain why we did not find conflict costs on obtained rewards as previously reported with the Simon task [35], which also involves externally-triggered conflict. As mentioned in the introduction, during the learning phase of that study [35], participants had to respond according to stimuli, some of which were associated with conflict. Thus, the fact that conflict was unavoidable might have increased its subjective cost. Furthermore, their design closely mirrored other effort discounting tasks, wherein people learn how much effort is needed to obtain a reward and, subsequently, show a preference for low effort options [3]. In contrast, in our study, conflict with distractors was fully orthogonal to the learning task. This allowed us to investigate how learning might be dynamically influenced by an unpredictable, and task-irrelevant, experience of conflict, rather than offering the opportunity to learn to predict upcoming conflict. Future work is clearly needed to investigate the conditions under which conflict might discount rewards, or be its effects may be cancelled out by other mechanisms.

**Fig S1.**
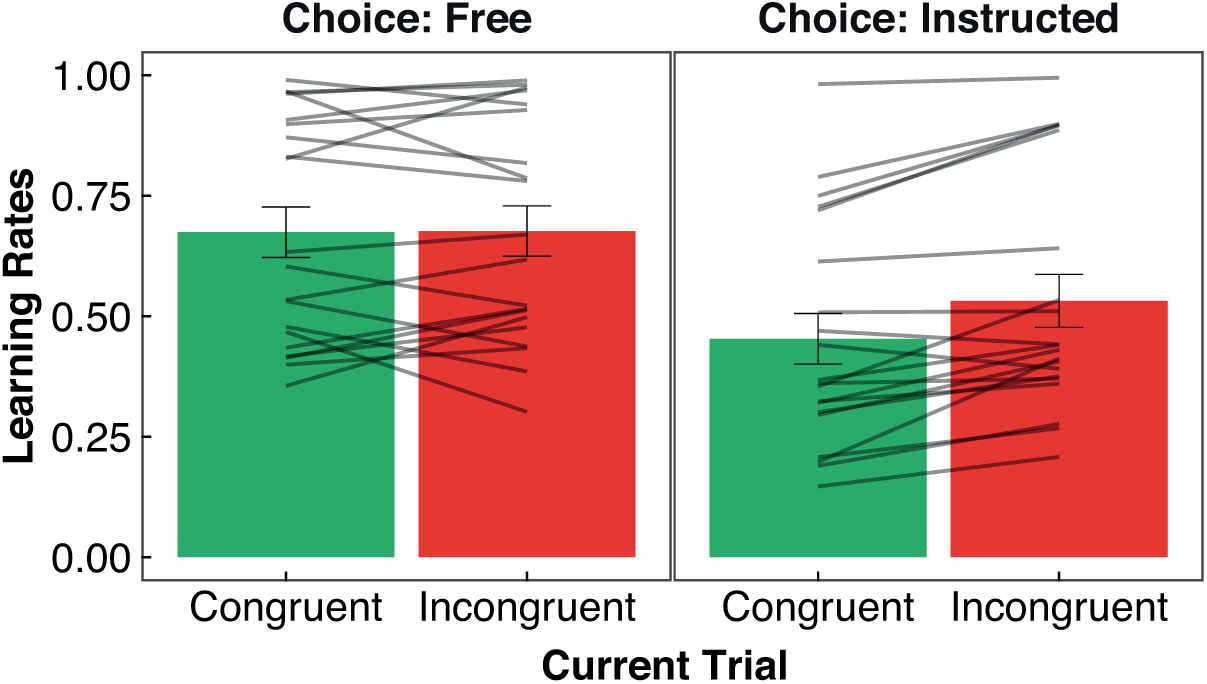
Estimated learning rates in m7. Learning rates (*α*) in m7 as a function of choice and current trial distractor-action congruency.

### S2 Text. Parameter vs. behaviour correlations

To investigate the relation between conflict avoidance and conflict adaptation effects, we assessed the relation between the estimated distractor bias parameter (as an index conflict avoidance) and conflict adaptation on RTs. Conflict adaptation effects were calculated as the difference between conflict effects (I *minus* C) for previously *congruent* minus previously incongruent trials. Thus, larger conflict adaptation reflects a greater reduction in conflict effects following incongruent trials. Since similar conflict adaptation was observed for free and instructed trials, we averaged over choice conditions. This analysis revealed a significant positive correlation between the distractor bias parameter, i.e. conflict avoidance, and conflict adaptation effects on RTs (see S2A Fig, Pearson’s correlation: *r* = 0.54, *t*_18_ = 2.72, *p* = .014). That is, participants who were better able to adapt their behaviour to reduce conflict costs on RTs were also more likely to avoid conflict when unnecessary (i.e. in the absence of strong value differences).

These results should be interpreted with care, given our relatively small sample size. Nonetheless, they suggest that participants’ sensitivity to conflict may be reflected in these two types of adaptive behaviours, rather than being a trade-off between them. It could have been hypothesised instead that participants who were worse at minimising RT costs would benefit most from avoiding conflict. Yet, this correlation implies that a common process of conflict monitoring and adaptation may underlie both types of behavioural responses. In fact, previous work has suggested that conflict signals can trigger both adjustments in cognitive control and conflict avoidance [[8,17,70] but see [19]].

Finally, it could have been hypothesised that having a larger choice bias might impair performance in the task, as participants choices might be too driven by the distractors rather than action values. Importantly, since the probably of left and right distractors was equate within each learning episode (i.e. between reversals), following the distractors’ suggestion would be equally likely to be helpful vs. unhelpful to task performance (i.e. 50/50 chance). Nevertheless, we tested this hypothesis by assessing the correlation between the estimated distractor bias parameter and average task performance, which showed no significant correlation (S2B Fig, Pearson’s correlation: *r* = -0.36, *t*_18_ = -1.63, *p* = .12). The independence of the distractor bias from effects on average performance was further corroborated by model simulations across a range of distractor bias parameters (S3 Fig).

**Fig S2.**
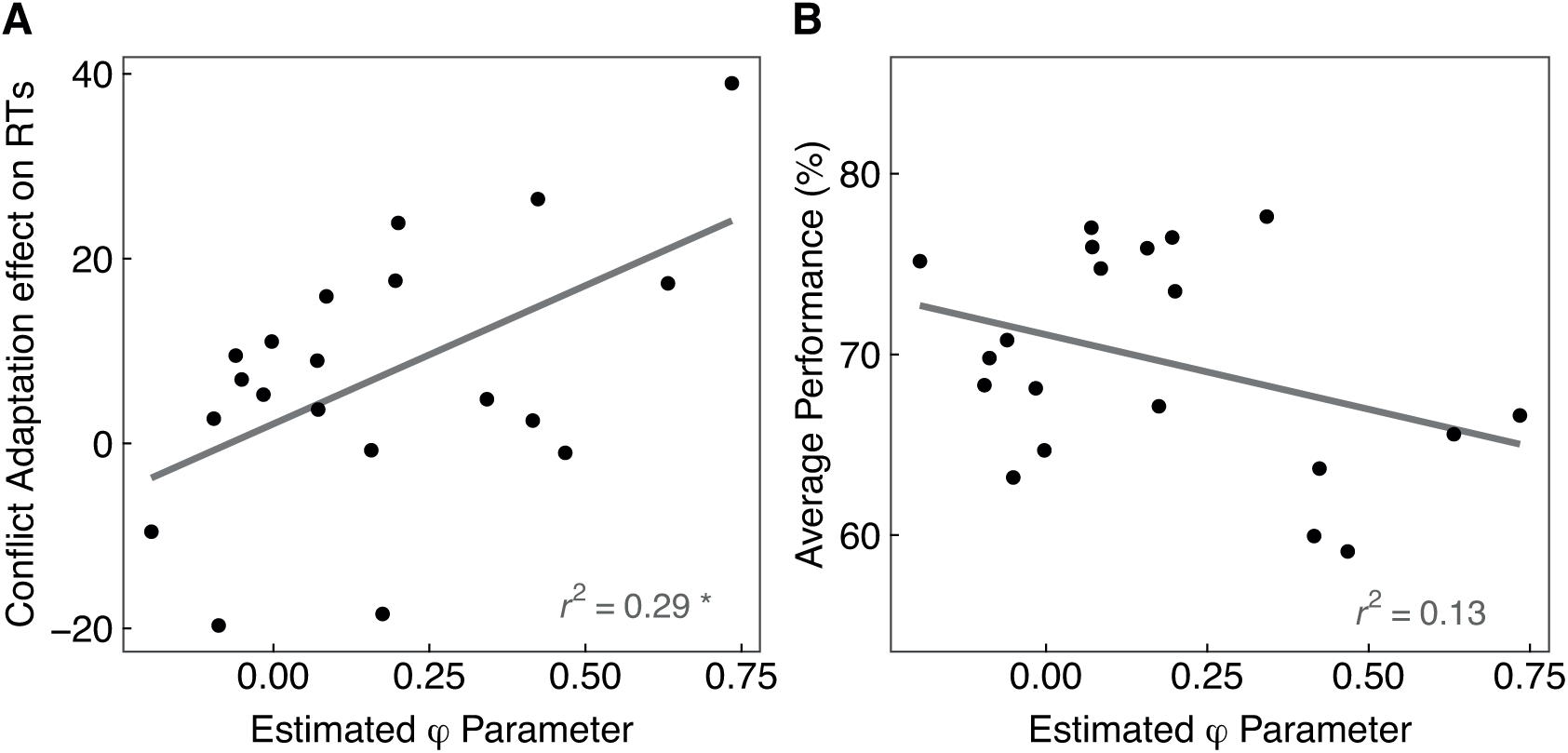
Parameter vs. behaviour correlations. Correlations between estimated distractor bias parameter and conflict adaptations effects on RTs (**A**), and average performance (**B**).

**Fig S3.**
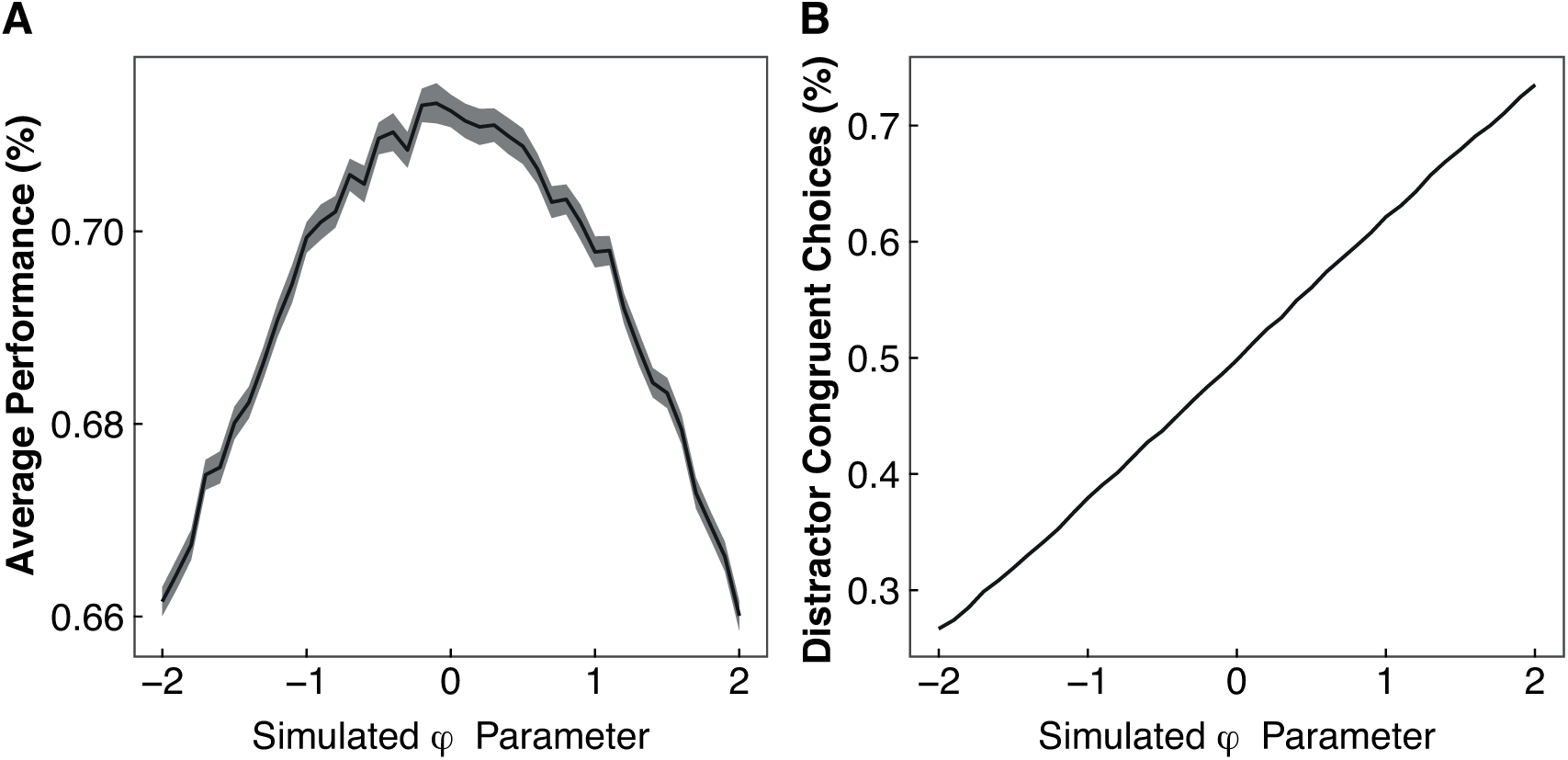
Parameter vs. simulated behaviour. Simulated behaviour in virtual datasets (*N* = 100) across a range of distractor bias (*φ*) values ([-2, 2], at intervals of 0.1; and constant *β* = 2, *α* = 0.6). **A.** Average performance. **B.** Percentage of distractor congruent choices. These findings show that, across this broad range of *φ* values (estimated *φ* varied less than 1), changes in average performance were minimal, whereas they were associated with very large differences in distractor bias on free choices (i.e. proportion of distractor congruent vs. incongruent choices).

### S3 Text. The critical role of the distractor bias parameter

To further test the need to consider an effect of distractors at the decision stage, we assessed whether an alternative model without that parameter was able to reproduce the behaviour observed in our participants. For this, we compared the generative performance of the winning model (m8) to an alternative model (m8_alt), which differed only in that the decision rule followed the standard softmax rule (i.e. without the added *φ*, equivalent to *φ* = 0; and with *α*_*Free*_ ≠ *α*_*Instructed_High Value*_ ≠ *α*_*Instructed_Low Value*_, as in in m8). First, we fitted this model to the real data. Next, we simulated used the average estimated parameter values to simulate data (*N* = 100). S4 Fig shows the simulated data from the alternative model, overlaid over the real data and simulations from the winning model (already displayed in Fig 3). This shows that the alternative model, without the distractor bias parameter, fails to qualitatively reproduce the observed distractor bias effect, i.e. higher proportion of distractor congruent than incongruent choices, from the start of learning episodes. This bias is instead clearly observed in simulations of the winning model (m8), which included the distractor bias parameter.

Therefore, we believe the poor generative performance of the alternative model, which fails to capture an important behavioural effect, is sufficient cause for rejecting it as a suitable model [71], hence excluding other models without a distractor bias from our model space.

**Fig S4.**
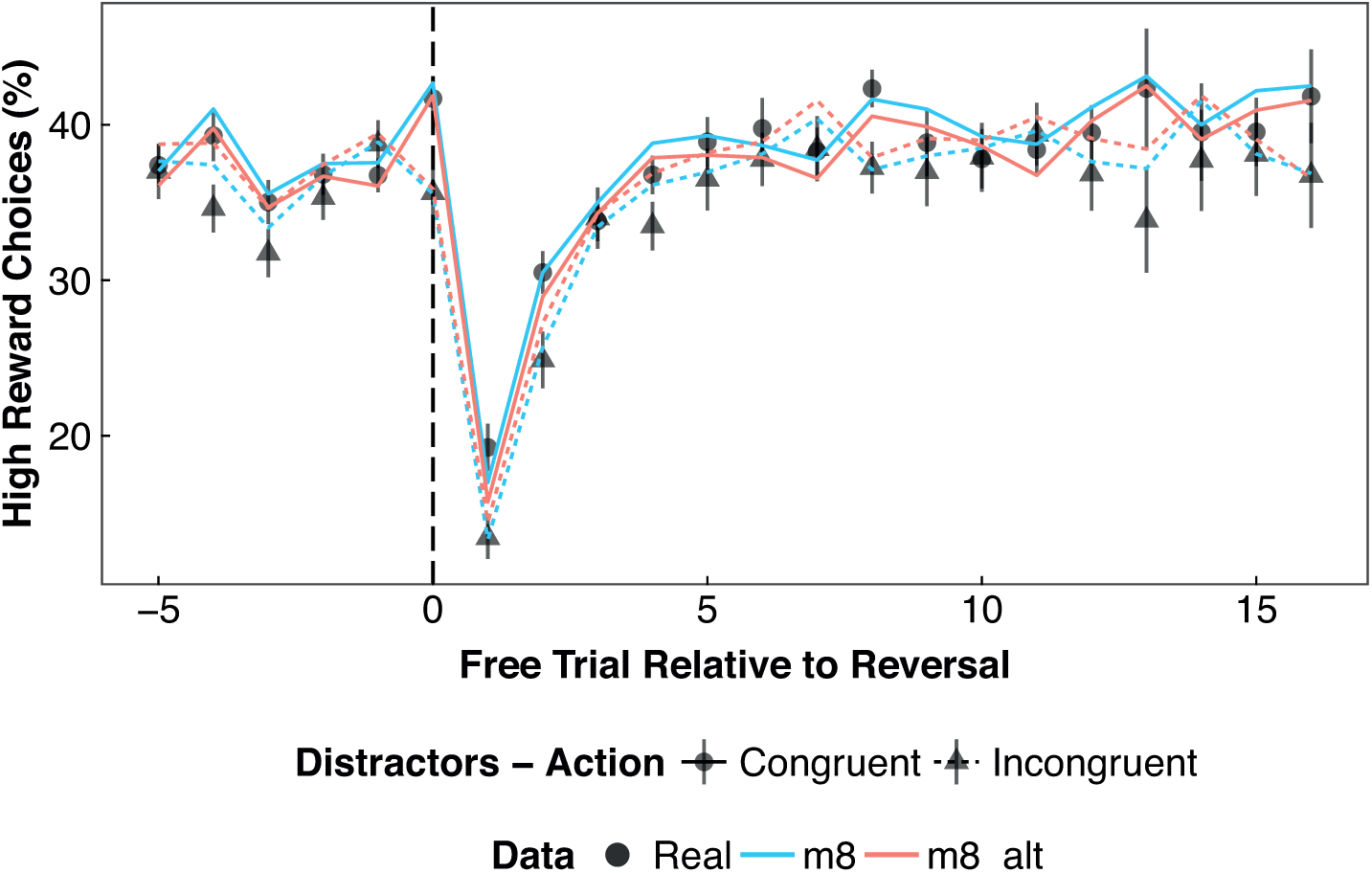
Model simulations. Simulations for the winning model (m8) and a similar model, without the distractor bias (*φ*) parameter (m8_alt).

### S4 Text. Parameter recovery: validation of the parameter optimisation procedure

To ensure that our task design and parameter optimisation procedure would indeed be able to identify the effects of conflict between action and distractors that we had hypothesised, we used simulated data based on pre-defined parameter values that should be recovered by the running the optimisation procedure on the simulated data. If our design or optimisation procedure were flawed, and would mistakenly introduce biases in the estimated parameters, then the recovered parameter values would differ from the parameters used in the “virtual participants”. We simulated virtual datasets (*N* = 100) based on 3 sets of parameter values. These aimed to capture 3 hypothetical results of action-distractor conflict: a) a cost of conflict on decision [*φ* = 0.2, *α*_*C*_ = *α*_*I*_ = 0.6], but no effect on learning; b) a cost of conflict on learning [*φ* = 0, *α*_*C*_ = 0.7, *α*_*I*_ = 0.5]; c) a benefit of conflict to learning [*φ* = 0, *α*_*C*_ = 0.5, *α*_*I*_ = 0.7]. We additionally tested parameter sets simulating effects on the decision and on learning simultaneously (d: [*φ* = 0.2, *α*_*C*_ = 0.7, *α*_*I*_ = 0.5]; e: [*φ* = 0.2, *α*_*C*_ = 0.5, *α*_*I*_ = 0.7]) These datasets were then submitted to the parameter optimisation procedure for the model that could assess an effect of conflict on the decision and on learning (i.e. m3 with the free parameters [*φ, α*_*C*_*, α*_*I*_]). The estimated parameters are displayed in S5 Fig. These results clearly show that the simulated parameter values were adequately recovered. Moreover, it shows that effects at the decision and learning stages are, at least theoretically, dissociable (as seen in S5 Fig, panels D. and E., where both effects were simulated). Consequently, the effects observed on our real participants (i.e. a cost of action-distractor conflict on the decision, but no effect on learning rates) cannot be attributed to our design or parameter estimation procedure introducing specific biases for or against finding particular effects.

**Fig S5.**
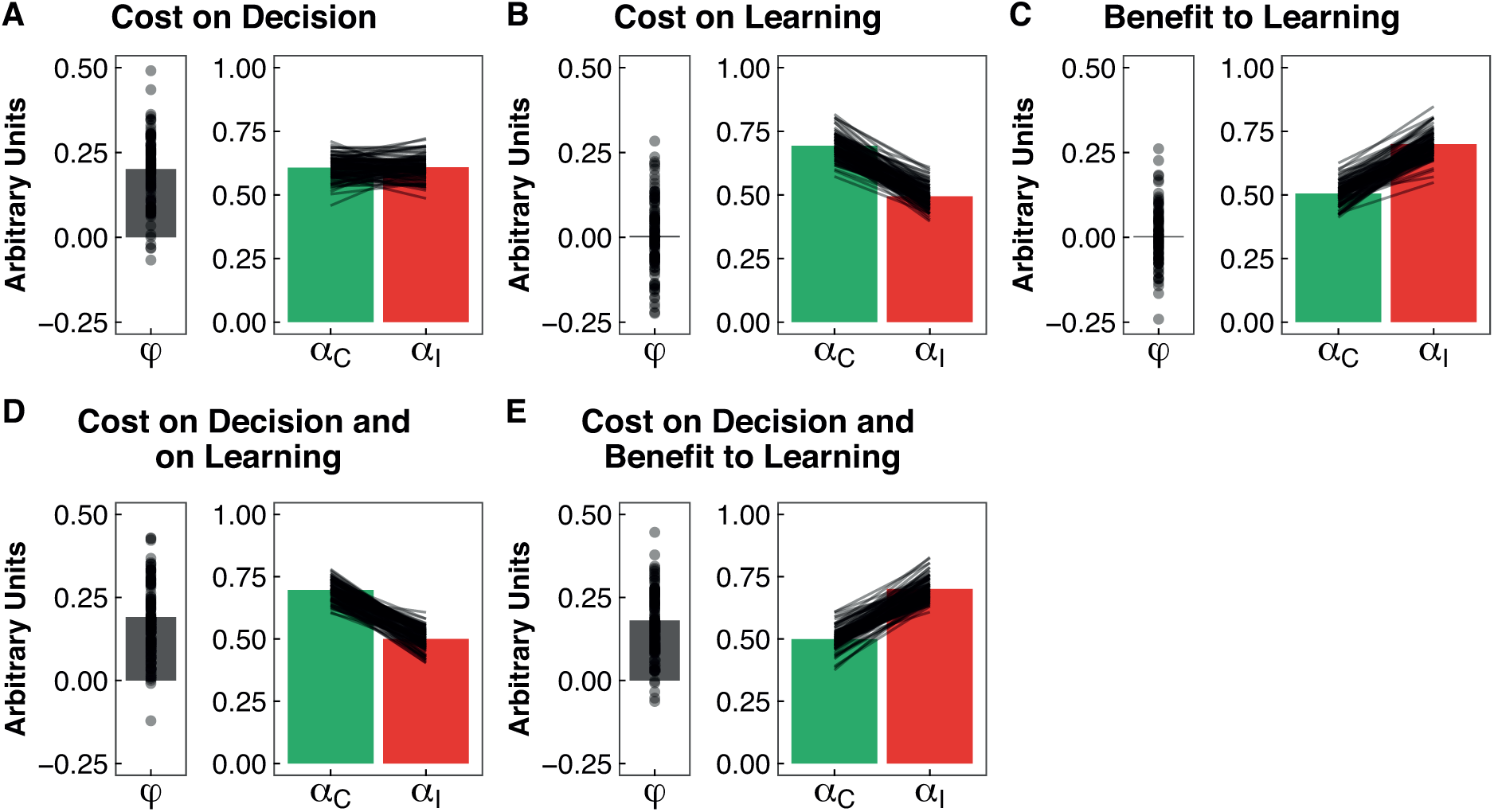
Parameter recovery. Validation of the parameter optimisation procedure, and ability of our model to capture dissociable effects of action-distractor conflict at the decision and learning stages. The bars display the average parameters obtained from the parameter optimisation procedure applied to simulated data based on three different parameter sets (*N* = 100, displayed as dots and lines). **A.** Results from data simulated with a cost of conflict on the decision [*φ* = 0.2, *α*_*C*_ = *α*_*I*_ = 0.6]. **B.** Results for data simulated with a cost of conflict on learning [*φ* = 0, *α*_*C*_ = 0.7, *α*_*I*_ = 0.5]. **C.** Results for data simulated with a benefit of conflict to learning [*φ* = 0, *α*_*C*_ = 0.5, *α*_*I*_ = 0.7]. Panels **D.** and **E.** combine the effect on the decision show in A, with the effects on learning in B and C, respectively. The results confirm that the simulated model parameters are adequately recovered by the optimisation procedure. C = Congruent, I = Incongruent.

## References

1. Skvortsova V, Palminteri S, Pessiglione M. Learning To Minimize Efforts versus Maximizing Rewards: Computational Principles and Neural Correlates. J Neurosci. 2014;34: 15621–15630. doi:10.1523/JNEUROSCI.1350-14.2014

2. Kurniawan IT, Seymour B, Talmi D, Yoshida W, Chater N, Dolan RJ. Choosing to Make an Effort: The Role of Striatum in Signaling Physical Effort of a Chosen Action. J Neurophysiol. 2010;104: 313–321. doi:10.1152/jn.00027.2010

3. Vassena E, Silvetti M, Boehler CN, Achten E, Fias W, Verguts T. Overlapping Neural Systems Represent Cognitive Effort and Reward Anticipation. PLoS ONE. 2014;9: e91008. doi:10.1371/journal.pone.0091008

4. Apps MAJ, Grima LL, Manohar S, Husain M. The role of cognitive effort in subjective reward devaluation and risky decision-making. Sci Rep. 2015;5: 16880. doi:10.1038/srep16880

5. Botvinick MM, Huffstetler S, McGuire JT. Effort discounting in human nucleus accumbens. Cogn Affect Behav Neurosci. 2009;9: 16–27. doi:10.3758/CABN.9.1.16

6. Kurniawan IT, Guitart-Masip M, Dayan P, Dolan RJ. Effort and Valuation in the Brain: The Effects of Anticipation and Execution. J Neurosci. 2013;33: 6160–6169. doi:10.1523/JNEUROSCI.4777-12.2013

7. Alexander WH, Brown JW. Computational Models of Performance Monitoring and Cognitive Control. Top Cogn Sci. 2010;2: 658–677. doi:10.1111/j.1756-8765.2010.01085.x

8. Botvinick MM. Conflict monitoring and decision making: Reconciling two perspectives on anterior cingulate function. Cogn Affect Behav Neurosci. 2007;7: 356–366. doi:10.3758/CABN.7.4.356

9. Janssens C, De Loof E, Boehler CN, Pourtois G, Verguts T. Occipital alpha power reveals fast attentional inhibition of incongruent distractors. Psychophysiology. 2017; doi:10.1111/psyp.13011

10. Scherbaum S, Fischer R, Dshemuchadse M, Goschke T. The dynamics of cognitive control: Evidence for within-trial conflict adaptation from frequency-tagged EEG. Psychophysiology. 2011;48: 591–600. doi:10.1111/j.1469-8986.2010.01137.x

11. Aben B, Verguts T, Van den Bussche E. Beyond trial-by-trial adaptation: A quantification of the time scale of cognitive control. J Exp Psychol Hum Percept Perform. 2017;43: 509–517. doi:10.1037/xhp0000324

12. Egner T. Congruency sequence effects and cognitive control. Cogn Affect Behav Neurosci. 2007;7: 380–390. doi:10.3758/CABN.7.4.380

13. Gratton G, Coles MG, Donchin E. Optimizing the use of information: strategic control of activation of responses. J Exp Psychol Gen. 1992;121: 480–506.

14. Botvinick MM, Braver T. Motivation and Cognitive Control: From Behavior to Neural Mechanism. Annu Rev Psychol. 2015;66: 83–113. doi:10.1146/annurev-psych-010814-015044

15. Braem S, King JA, Korb FM, Krebs RM, Notebaert W, Egner T. The Role of Anterior Cingulate Cortex in the Affective Evaluation of Conflict. J Cogn Neurosci. 2016;29: 137–149. doi:10.1162/jocn_a_01023

16. Dreisbach G, Fischer R. Conflicts as aversive signals. Brain Cogn. 2012;78: 94–98. doi:10.1016/j.bandc.2011.12.003

17. Dignath D, Kiesel A, Eder AB. Flexible conflict management: Conflict avoidance and conflict adjustment in reactive cognitive control. J Exp Psychol Learn Mem Cogn. 2015;41: 975–988. doi:10.1037/xlm0000089

18. Schouppe N, Demanet J, Boehler CN, Ridderinkhof KR, Notebaert W. The Role of the Striatum in Effort-Based Decision-Making in the Absence of Reward. J Neurosci. 2014;34: 2148–2154. doi:10.1523/JNEUROSCI.1214-13.2014

19. Schouppe N, Ridderinkhof KR, Verguts T, Notebaert W. Context-specific control and context selection in conflict tasks. Acta Psychol (Amst). 2014;146: 63–66. doi:10.1016/j.actpsy.2013.11.010

20. Desender K, Buc Calderon C, Van Opstal F, Van den Bussche E. Avoiding the conflict: Metacognitive awareness drives the selection of low-demand contexts. J Exp Psychol Hum Percept Perform. 2017;43: 1397–1410. doi:10.1037/xhp0000391

21. Olson JA, Amlani AA, Raz A, Rensink RA. Influencing choice without awareness. Conscious Cogn. 2015; doi:10.1016/j.concog.2015.01.004

22. Sidarus N, Haggard P. Difficult action decisions reduce the sense of agency: A study using the Eriksen flanker task. Acta Psychol (Amst). 2016;166: 1–11. doi:10.1016/j.actpsy.2016.03.003

23. Kiesel A, Wagener A, Kunde W, Hoffmann J, Fallgatter AJ, Stöcker C. Unconscious manipulation of free choice in humans. Conscious Cogn. 2006;15: 397–408. doi:10.1016/j.concog.2005.10.002

24. Klapp ST, Hinkley LB. The negative compatibility effect: Unconscious inhibition influences reaction time and response selection. J Exp Psychol Gen. 2002;131: 255–269. doi:10.1037/0096-3445.131.2.255

25. O’Connor PA, Neill WT. Does subliminal priming of free response choices depend on task set or automatic response activation? Conscious Cogn. 2011;20: 280–287. doi:10.1016/j.concog.2010.08.007

26. Schlaghecken F, Eimer M. Masked prime stimuli can bias “free” choices between response alternatives. Psychon Bull Rev. 2004;11: 463–468. doi:10.3758/BF03196596

27. Sidarus N, Vuorre M, Haggard P. How action selection influences the sense of agency: An ERP study. Neuroimage. 2017;150: 1–13. doi:10.1016/j.neuroimage.2017.02.015

28. euchies M, Demanet J, Sidarus N, Haggard P, Stevens MA, Brass M. Influences of unconscious priming on voluntary actions: Role of the rostral cingulate zone. Neuroimage. 2016;135: 243–252. doi:10.1016/j.neuroimage.2016.04.036

29. Frank MJ, Gagne C, Nyhus E, Masters S, Wiecki TV, Cavanagh JF, et al. fMRI and EEG Predictors of Dynamic Decision Parameters during Human Reinforcement Learning. J Neurosci. 2015;35: 485–494. doi:10.1523/JNEUROSCI.2036-14.2015

30. Shenhav A, Musslick S, Lieder F, Kool W, Griffiths TL, Cohen JD, et al. Toward a Rational and Mechanistic Account of Mental Effort. Annu Rev Neurosci. 2017;40: 99–124. doi:10.1146/annurev-neuro-072116-031526

31. Fritz J, Dreisbach G. Conflicts as aversive signals: Conflict priming increases negative judgments for neutral stimuli. Cogn Affect Behav Neurosci. 2013;13: 311–317. doi:10.3758/s13415-012-0147-1

32. Fritz J, Dreisbach G. The Time Course of the Aversive Conflict Signal. Exp Psychol. 2015;62: 30–39. doi:10.1027/1618-3169/a000271

33. Wenke D, Fleming SM, Haggard P. Subliminal priming of actions influences sense of control over effects of action. Cognition. 2010;115: 26–38. doi:10.1016/j.cognition.2009.10.016

34. Chambon V, Sidarus N, Haggard P. From action intentions to action effects: how does the sense of agency come about? Front Hum Neurosci. 2014;8: 320. doi:10.3389/fnhum.2014.00320

35. Cavanagh JF, Masters SE, Bath K, Frank MJ. Conflict acts as an implicit cost in reinforcement learning. Nat Commun. 2014;5: comms6394. doi:10.1038/ncomms6394

36. Simon JR, Rudell AP. Auditory S-R compatibility: The effect of an irrelevant cue on information processing. J Appl Psychol. 1967;51: 300–304. doi:10.1037/h0020586

37. Guitart-Masip M, Huys QJM, Fuentemilla L, Dayan P, Duzel E, Dolan RJ. Go and no-go learning in reward and punishment: Interactions between affect and effect. Neuroimage. 2012;62: 154–166. doi:10.1016/j.neuroimage.2012.04.024

38. Swart JC, Froböse MI, Cook JL, Geurts DE, Frank MJ, Cools R, et al. Catecholaminergic challenge uncovers distinct Pavlovian and instrumental mechanisms of motivated (in)action. eLife. 2017;6: e22169. doi:10.7554/eLife.22169

39. Sutton RS, Barto AG. Reinforcement Learning: An Introduction. Cambridge, MA: MIT Press; 1998.

40. Watkins CJCH, Dayan P. Q-learning. Mach Learn. 1992;8: 279–292. doi:10.1007/BF00992698

41. Voss M, Chambon V, Wenke D, Kühn S, Haggard P. In and out of control: brain mechanisms linking fluency of action selection to self-agency in patients with schizophrenia. Brain. 2017;140: 2226–2239. doi:10.1093/brain/awx136

42. Kool W, McGuire JT, Rosen ZB, Botvinick MM. Decision Making and the Avoidance of Cognitive Demand. J Exp Psychol Gen. 2010;139: 665–682. doi:10.1037/a0020198

43. Kool W, Botvinick M. The intrinsic cost of cognitive control. Behav Brain Sci. 2013;36: 697–698. doi:10.1017/S0140525X1300109X

44. Shenhav A, Botvinick MM, Cohen JD. The Expected Value of Control: An Integrative Theory of Anterior Cingulate Cortex Function. Neuron. 2013;79: 217–240. doi:10.1016/j.neuron.2013.07.007

45. Mulder MJ, van Maanen L, Forstmann BU. Perceptual decision neurosciences – A model-based review. Neuroscience. 2014;277: 872–884. doi:10.1016/j.neuroscience.2014.07.031

46. Ratcliff R, Smith PL, Brown SD, McKoon G. Diffusion Decision Model: Current Issues and History. Trends Cogn Sci. 2016;20: 260–281. doi:10.1016/j.tics.2016.01.007

47. Hanks TD, Mazurek ME, Kiani R, Hopp E, Shadlen MN. Elapsed Decision Time Affects the Weighting of Prior Probability in a Perceptual Decision Task. J Neurosci. 2011;31: 6339–6352. doi:10.1523/JNEUROSCI.5613-10.2011

48. O’Doherty J, Dayan P, Schultz J, Deichmann R, Friston K, Dolan RJ. Dissociable Roles of Ventral and Dorsal Striatum in Instrumental Conditioning. Science. 2004;304: 452–454. doi:10.1126/science.1094285

49. Kool W, Getz SJ, Botvinick MM. Neural Representation of Reward Probability: Evidence from the Illusion of Control. J Cogn Neurosci. 2013;25: 852–861. doi:10.1162/jocn_a_00369

50. Leotti LA, Delgado MR. The Inherent Reward of Choice. Psychol Sci. 2011;22: 1310–1318. doi:10.1177/0956797611417005

51. Langer EJ. The illusion of control. J Pers Soc Psychol. 1975;32: 311–328. doi:10.1037/0022-3514.32.2.311

52. Lefebvre G, Lebreton M, Meyniel F, Bourgeois-Gironde S, Palminteri S. Behavioural and neural characterization of optimistic reinforcement learning. Nat Hum Behav. 2017;1: 0067. doi:10.1038/s41562-017-0067

53. Sharot T. The optimism bias. Curr Biol. 2011;21: R941-945. doi:10.1016/j.cub.2011.10.030

54. Kornell N, Metcalfe J. Study efficacy and the region of proximal learning framework. J Exp Psychol Learn Mem Cogn. 2006;32: 609–622. doi:10.1037/0278-7393.32.3.609

55. Murty VP, DuBrow S, Davachi L. The Simple Act of Choosing Influences Declarative Memory. J Neurosci. 2015;35: 6255–6264. doi:10.1523/JNEUROSCI.4181-14.2015

56. Gureckis TM, Markant DB. Self-Directed Learning: A Cognitive and Computational Perspective. Perspect Psychol Sci. 2012;7: 464–481. doi:10.1177/1745691612454304

57. Bellebaum C, Jokisch D, Gizewski ER, Forsting M, Daum I. The neural coding of expected and unexpected monetary performance outcomes: Dissociations between active and observational learning. Behav Brain Res. 2012;227: 241–251. doi:10.1016/j.bbr.2011.10.042

58. Kobza S, Bellebaum C. Processing of action-but not stimulus-related prediction errors differs between active and observational feedback learning. Neuropsychologia. 2015;66: 75–87. doi:10.1016/j.neuropsychologia.2014.10.036

59. Cohen MX, Cavanagh JF. Single-Trial Regression Elucidates the Role of Prefrontal Theta Oscillations in Response Conflict. Front Psychol. 2011;2. doi:10.3389/fpsyg.2011.00030

60. Cavanagh JF, Eisenberg I, Guitart-Masip M, Huys Q, Frank MJ. Frontal Theta Overrides Pavlovian Learning Biases. J Neurosci. 2013;33: 8541–8548. doi:10.1523/JNEUROSCI.5754-12.2013

61. Brainard DH. The Psychophysics Toolbox. Spat Vis. 1997;10: 433–436.

62. Kleiner M, Brainard DH, Pelli DG. What’s new in Psychtoolbox-3? Perception. 2007;36. doi:10.1068/v070821

63. Pelli DG. The VideoToolbox software for visual psychophysics: transforming numbers into movies. Spat Vis. 1997;10: 437–442.

64. Schwarz G. Estimating the Dimension of a Model. Ann Stat. 1978;6: 461–464. doi:10.1214/aos/1176344136

65. Daunizeau J, Adam V, Rigoux L. VBA: A Probabilistic Treatment of Nonlinear Models for Neurobiological and Behavioural Data. PLoS Comput Biol. 2014;10: e1003441. doi:10.1371/journal.pcbi.1003441

66. Stephan KE, Penny WD, Daunizeau J, Moran RJ, Friston KJ. Bayesian Model Selection for Group Studies. Neuroimage. 2009;46: 1004–1017. doi:10.1016/j.neuroimage.2009.03.025

67. Daw ND, Gershman SJ, Seymour B, Dayan P, Dolan RJ. Model-Based Influences on Humans’ Choices and Striatal Prediction Errors. Neuron. 2011;69: 1204–1215. doi:10.1016/j.neuron.2011.02.027

68. Palminteri S, Lefebvre G, Kilford EJ, Blakemore S-J. Confirmation bias in human reinforcement learning: Evidence from counterfactual feedback processing. PLoS Comput Biol. 2017;13. doi:10.1371/journal.pcbi.1005684

69. Schouppe N, Braem S, Houwer JD, Silvetti M, Verguts T, Ridderinkhof KR, et al. No pain, no gain: the affective valence of congruency conditions changes following a successful response. Cogn Affect Behav Neurosci. 2015;15: 251–261. doi:10.3758/s13415-014-0318-3

70. Dreisbach G, Fischer R. If it’s hard to read… try harder! Processing fluency as signal for effort adjustments. Psychol Res. 2011;75: 376–383. doi:10.1007/s00426-010-0319-y

71. Palminteri S, Wyart V, Koechlin E. The Importance of Falsification in Computational Cognitive Modeling. Trends Cogn Sci. 2017; doi:10.1016/j.tics.2017.03.011

